# Climate stress priming of juvenile Southern California giant kelp (*Macrocystis pyrifera*) to thermal extremes

**DOI:** 10.64898/2026.01.05.697421

**Authors:** Lauren L. Smith, Briana Le, Skye T. Krainer, Andie McNeil, Abigail P. Van Slyke, Halley E. Froehlich

## Abstract

Climate change threatens food production across the globe, creating challenges for food systems. Aquaculture, including seaweed production, is expanding while being threatened by global climate stressors, including increasing extreme events. Marine aquaculture is especially vulnerable to heatwaves, which can rapidly raise temperatures above the physiological limits of some organisms. While several interventions to increase resilience to climate impacts are being explored, ‘priming’ has emerged as a possible adaptation for seaweeds that maintains genetic diversity but hardens individuals to stressors later in life. California has a developing seaweed sector while also experiencing some of the most extreme marine heatwave conditions on record. We explore temperature impacts and priming – exposing an earlier stage of an organism to a mild stressor to prepare the individual for future stress – on giant kelp *Macrocystis pyrifera*, an important foundation species along the West Coast of the United States. Our experiments focused on the juvenile sporophyte stage on miniaturized spools, from approximately one week before outplanting size to one week after. First, we determined the reaction norms of *M. pyrifera* in waters ranging from 5 to 30°C at the outplanting stage. Next, we explored how priming (heat + or – nutrients) in a hatchery setting prepares *M. pyrifera* for outplanting to a marine heatwave. To assess experimental outcomes, we took measures of growth, survival, photosynthetic function (*Fv/Fm*), and carbon and nitrogen assimilation via isotopes. We found temperatures above 20°C had significant negative impacts on all metrics of performance during the juvenile sporophyte stage. Further, we determined heat priming in conjunction with hatchery level (+) nutrients resulted in overall increased performance when exposed to a marine heatwave. These findings support the continued exploration of priming as a tool for climate resilience and can inform current hatchery practices for aquaculture practitioners looking to improve crop outcomes for this species.

## INTRODUCTION

Food production and security are increasingly threatened by climate change and its associated extreme variability (1,2). A shifting climate can negatively impact food systems (e.g., crop yield and/or quality (3)) through average changes in conditions (e.g., mean temperature increase), but increasingly, extreme events (e.g., atmospheric rivers, heatwaves) (4) are creating unprecedented acute challenges to wild and farmed systems alike (5,6). For example, marine heatwaves (MHWs) – an acute to prolonged period of unusually high temperatures (7), a type of ‘environmental whiplash’ (8) – change thermal conditions so rapidly it can surpass the physiological ability of some organisms to sufficiently respond (9). And while the social and ecological impacts of extremes and possible adaptations are increasingly studied for certain for sectors (e.g., terrestrial agriculture), aquaculture (i.e., aquatic farming) has been comparatively overlooked (10).

Total aquaculture production has exceeded all wild capture fisheries landings, playing an increasingly important role in the global food system (11), but is also under threat from global climate change (12–15). Aquaculture includes production of at least 730 species or taxa globally, spanning finfish, invertebrates, and seaweed across freshwater, brackish, and marine environments (16). Given the complexities of the systems in aquatic farming, pressures span ocean acidification, sea level rise, flooding, and marine heatwaves, to name a few (13). Saltwater aquaculture is particularly vulnerable to marine heatwaves, but the observations and science is just emerging (2,10). The dearth of climate impact and adaptation knowledge is particularly stark for farmed seaweeds; a sector which accounts for 36 million tonnes of wet weight ever year , $17 billion dollars (USD) globally (17), and is a rising sector in numerous western countries, including the United States (18). At least 32 species of algae are reportedly farmed worldwide (16) and 40% of these belong to the class Phaeophyceae – also known as brown algae (14). Kelps are particularly important subset of browns, used as fertilizers, feed, and biofuels, and are seen as a potential future, low carbon food source for humans (14).

While there are several on-farm, organismal interventions to help adapt to climate change and potential extremes (e.g., genetic selection) (10), one emerging adaptation for species lacking a nervous system, such as seaweeds, is ‘*priming*’ (15). In scientific literature, priming is also referred to as ‘conditioning’, ‘hardening’, and even ‘acclimation’ (19). Used in land-based crop agriculture (20–22), and also being tested on invertebrates (23), priming occurs when an organism is exposed to a mild stressor at a critical point in their life stage in a hatchery, resulting in a ‘stress memory’, which prepares (i.e., primes) the individual for futures stressful conditions (19,24). The stress memory is defined as a plastic change at the phenotypic level, induced by a non-lethal stressor that does not change the underlying genetics for the organism (19,24). Although currently unknown for most seaweeds, mechanisms of priming in other organisms include epigenetic modifications, such as microRNA, histone modifications, and DNA methylation, and resulting changes to gene expression, physiology, and metabolism that can be retained over short- (i.e., days to weeks) or long-term timeframes (i.e., lifetime, generations if bred) (19,24). It is also hypothesized that priming needs to occur on earlier stages of seaweed and marine plants to be most effective due to cellular changes in growth and development (19). Importantly, in some regions (e.g., USA), adaptative interventions for new or non-established species, such as farmed seaweeds, are required to protect surrounding genetic diversity, thus limiting more typical selective practices. For example, Alaska’s ‘50-50 rule’ – being adopted by other states, including California – where spore collection must be from more than 50 individuals within 50km of the outplanting site (25). While MHW intensity, frequency, and extent increase globally (26), considerable impacts and research across the California current provide an ideal region to explore such impacts and interventions.

The state of California has not only seen some of the most extreme MHWs, impacting socio-ecological ecosystems across the coast (27–29), there is also an emerging seaweed sector. The 2014-2016 MHW off the west coast of North America, known as “the blob”, had devastating, although varied, impacts on wild populations of giant kelp, *Macrocystis pyrifera* (30). *M. pyrifera* is a foundation species (31,32), playing a large role in nearshore community structure and function (33) and supporting over 800 other species off the California coast (34). In 2020, Ocean Rainforest broke ground on the first offshore pilot farm (4.4nm from shore) in California, growing *M. pyrifera*. Yet, recent climate studies project a potential decrease in suitability for seaweed culture across the West Coast of the United States, with southern California being particularly vulnerable (35). Certain life stages, such as adult sporophytes and gametophytes, of these species have received a lot of research attention (36–40). In fact, a recent study found heat priming gametophytes for four weeks led to better outcomes for their sporophytes when later exposed to high temperatures (41). However juvenile sporophytes – the stage farms outplant into the wild – are understudied. A recent study did find MHW intensity and duration negatively affect growth in young sporophytes (42), but the stage (older juveniles) and context (non-spooled) of the study were not aquaculture specific. This provides an opportune moment to partner with farms to explore interventions at the hatchery stage that could improve growth and survival during the outplant stage at higher temperatures.

The objective of our research was to 1) assess how temperature impacts growth at this early sporophyte stage and 2) explore how priming in the hatchery stage may trigger shifts in morphology and/or physiology that lead to an increased ability to withstand marine heatwaves in the future at the individual (singe blade) and population scale (spool). These include measures of growth, survival, photosynthetic function, and carbon and nitrogen assimilation (via stable isotopes). First, we manipulated the temperature for early juvenile *M. pyrifera* once they reached outplanting size to determine how temperatures between 5 and 30°C will impact kelp growth. Next, we manipulated the available nutrients and temperature for early juvenile *M. pyrifera* sporophytes during short-term priming that took place during the hatchery stage. Following a recovery period, the sporophytes underwent a simulated outplanting into a MHW to determine the efficacy of priming in preparing sporophytes for outplanting.

Understanding the viability of heat priming sporophytes at this early stage will inform the active measures kelp farms can take to climate proof their crop as human-induced climate change continues.

## MATERIALS AND METHODS

### 2.1 Spore collection and culture

*M. pyrifera* spore collection and culture were provided by Ocean Rainforest. All sorus material was collected via SCUBA between 7 and 12 meters of water within 50km of the farm, located just north of Santa Barbara and 4.4 nautical miles off the coast. Their internal protocols were followed for sporulation.

To simulate growing conditions of a typical kelp hatchery, ‘mini-spools’ were co-designed (N = 72 total number of spools) with Ocean Rainforest to emulate the structure and function of a seeded spool (a full-sized spool is 15” x 2” sterilized PVC pipe). Specifically, six centimeters of proprietary twine was tightly wound around 4” x 1” PVC to create the mini-spools for *M. pyrifera* seeding (Fig 1). At Ocean Rainforest, the mini-spools were submerged in a gametophyte solution (40,000 spores/mL) for 24 hours and then maintained in their facility until transitioning to the sporophyte life stage. This size was selected as it is hypothesized that priming needs to occur in the earlier stages of seaweeds to be most effective due to cellular changes in growth and development (i.e., ‘reset’) (19) (Fig 1). One important note, based on logistical challenges and labor capacity at Ocean Rainforest, there were some slight differences in mini-spool creation: in the priming experiment phase 1, two spools were raised in the same 3L mesocosm jars (n = 12 jars), while in phase 2 (n = 24) mini-spools were reared together in a single, large aquarium tank. The mini-spools for the kelp performance experiment mirrored phase 2. Regardless, once sporophytes were visible (<1mm; approximately 1-week before outplanting size), the mini-spools were transported in a single cooler on ice to UC Santa Barbara for the experiments. During an approximate 1-hour window of time, mini-spools were carefully and individually wrapped in plastic wrap to maintain moisture, placed on ice in a single cooler while transported, then kept refrigerated at 10°C until placed into their designated tanks for initial acclimation that same day.

**Fig 1.**
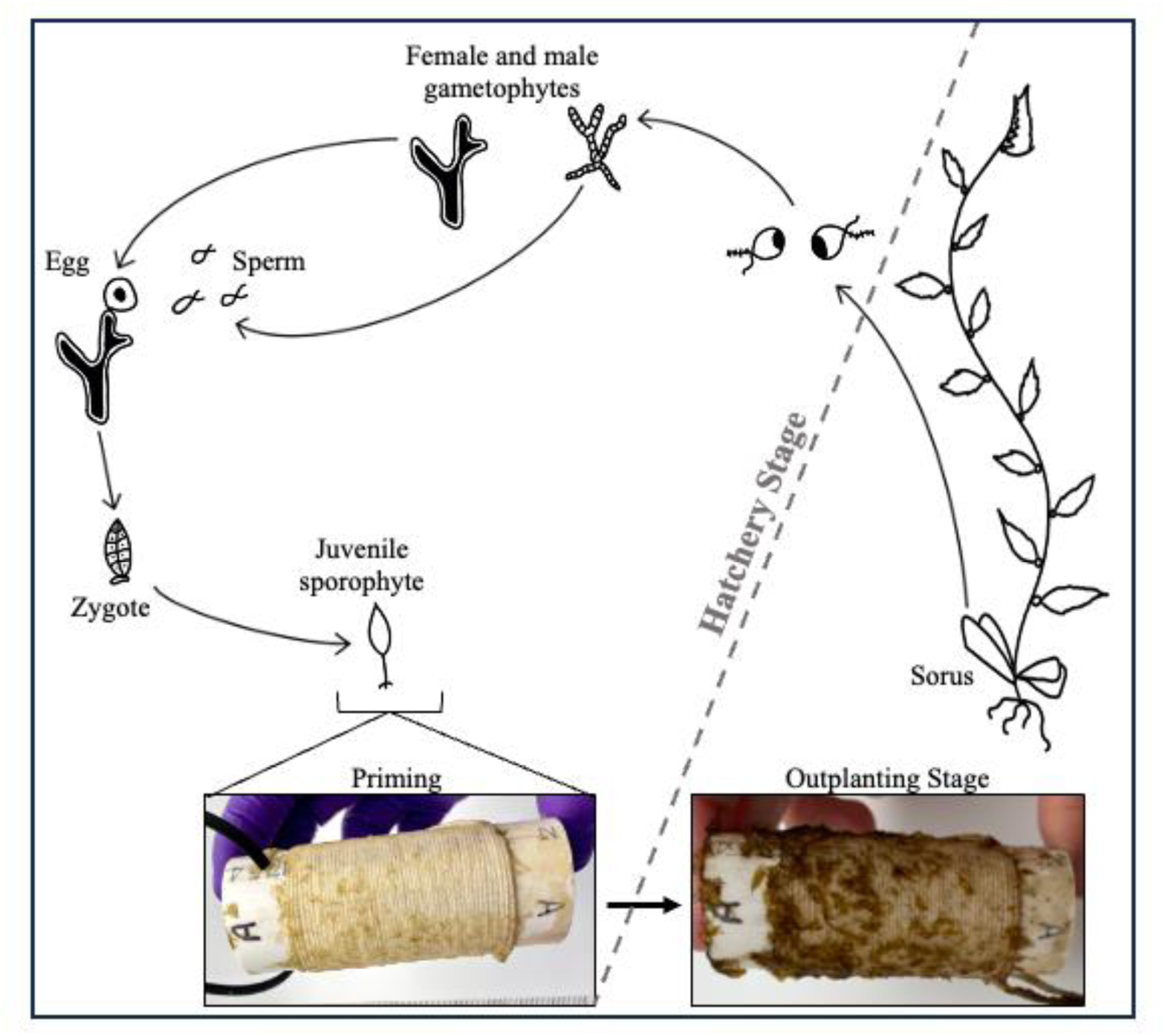
The life cycle of *Macrocystis pyrifera*. *Macrocystis pyrifera* has a heteromorphic alternation of generations. Images at the bottom of the life cycle show the giant kelp growth on spools in the hatchery stage where priming occurs and at the conclusion of the experiment.

### 2.2 Experimental Conditions

The kelp performance and priming experiments took place in the Froehlich Lab Aqualogic tank system at UC Santa Barbara. The tank system was equipped with six headers to control temperatures and 12 experimental tanks (Fig 2). Header tanks were connected to chilled seawater (10°C), which gravity fed into the experimental tanks, acting as a water bath for the 3L mesocosm jars (two per tank). After transport from Ocean Rainforest, mini spools were hung from the top of each mesocosm jar (kelp performance N = 24 and priming N = 48). Inside each jar, we placed UV-treated aquarium rocks to maintain negative buoyancy. Above each tank was a 12-inch grow light (Barriana T5 LED full spectrum grow light; PAR = 50 in tanks with no water) with a light-cycle of 14:10 day to night, replicating Ocean Rainforest’s protocol.

**Fig 2.**
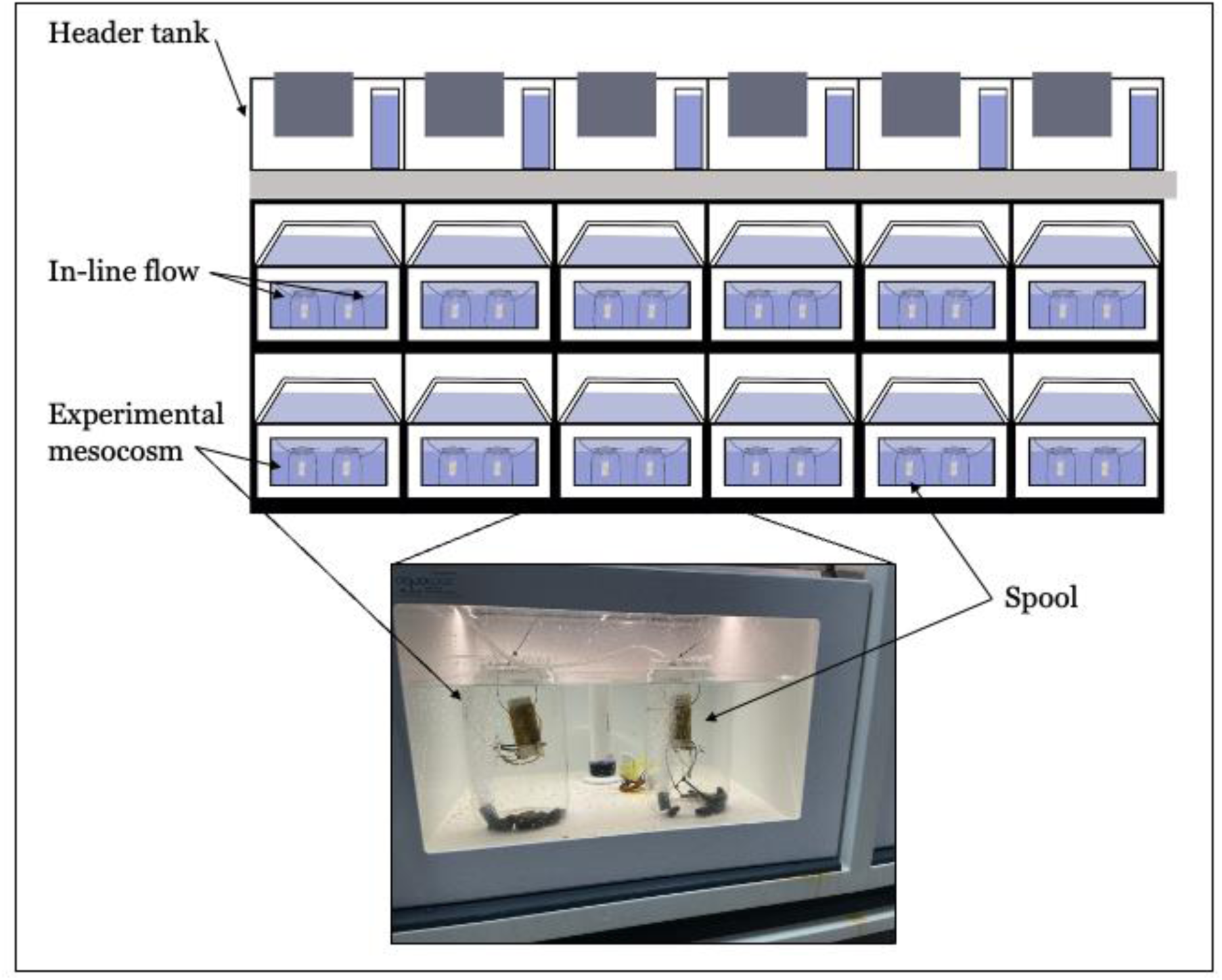
Tank schematic for experimental set up. Graphic depicting the experimental set-up where six temperature-controlled header tanks gravity feed into 12 experimental tanks. Each experimental tank is equipped with two closed mesocosms. The photograph shows the spools hanging from the mesocosms inside an experimental tank.

Both experiments were broken into a hatchery and outplanting stage, described in more detail below. During the hatchery stage (7-days), the seawater was exposed to a 4-hour UV treatment to reduce bacteria and diatom contamination. Nutrients were added for the hatchery stage using F2, a concentrated Guillard f/2 formula produced by the Mercer of Montana and used inhouse by Ocean Rainforest. For the outplanting stage (10-days) of the experiment, all mini-spools were flipped to hang from the opposite side, and the water in each mesocosm was exchanged for non-UV treated seawater, with no additional nutrients added. The latter, non-treated water was meant to simulate ‘real world’ ocean conditions.

Temperatures across both experiments were tracked using a mix of Onset HOBO pendant temperature and temperature/light loggers. A total of 24 loggers were deployed, two in each experimental tank, that logged temperature (°C) every 30 minutes. For each treatment, these temperatures were averaged during each discrete temperature window.

### 2.3 Kelp performance curve

To help inform and interpret the priming experiment, we conducted a performance curve experiment on *M. pyrifera* juvenile sporophytes across a thermal gradient (N = 24 mini-spools; n = 4 per thermal treatment) from October 7^th^ through 24^th^, 2024. In order to test the comparable outplant stage of the priming experiment, all mini-spools were grown at 10°C with hatchery levels of nutrients (1mL of F2 per L of seawater) for seven days in the lab (i.e., hatchery stage). Once the hatchery stage was complete, headers were set to temperatures between 5 and 30°C, in five-degree increments (i.e., outplanting stage). Mini-spools were exposed to these temperatures for seven days before all headers were returned to 10°C for two days of recovery and subsequent data collection.

### 2.4 Priming giant kelp

To test the performance of primed *M. pyrifera*, we conducted a 2-factor fully crossed experiment during the hatchery stage, split across two phases (N = 48; 24 per phase). Given the known importance and potential interactions between temperature and nutrients (39,43), we explicitly accounted for the potential interactive effects of the ‘nutrient rescue hypothesis’ in the experiment design. The nutrient rescue hypothesis posits that supplemental nutrients allows organisms to withstand greater environmental stressors, such as increased temperatures (44,45). The treatments included a control (no priming; n = 8), and three different priming treatments. All treatments occurred during the 7-day hatchery stage. The first priming factor was ‘nutrient priming’, where half the mini-spools were given ‘high’ nutrient concentrations (1mL of F/2 L^-1^), while the others were given half (0.5mL of F/2 L^-1^). Note, the high nutrient treatment is the amount typically administered in the hatchery. The second priming factor was ‘heat priming’, a short-term, moderate thermal heat exposure. The heat priming consisted of 48 hours at higher temperatures, beginning after a 2-day acclimation period. Tanks were raised to 12°C for 24 hours and raised again to 15°C for 24 hours before being returned to 10°C for two days of recovery. This short exposure regiment aligns with the current standing priming science for macroalgae (19). Temperatures were based on thermal dynamics near the farm (NSF LTER, NOAA buoy, and Ocean Rainforest hobo logger) and known thermal bounds of adult sporophytes (39). Phase 1 was conducted from March 22^nd^ to April 2^nd^, 2024 and included the fully primed (nutrients and heat combined) and control treatments. Phase 2 was conducted from July 12^th^ to 29^th^, 2024 and included the nutrient primed and heat primed only treatments.

After seven days in the hatchery stage, all mini-spools experienced a simulated outplanting – although the twine remained spooled – into a MHW (i.e., no added F/2 nutrients, and no-UV treated seawater). This began by raising all header temperatures to 12°C for 24 hours, then to 15°C for another 24 hours. Finally, the temperature was raised to 18°C for five days before a 2-day recovery period (Fig 3).

**Fig 3.**
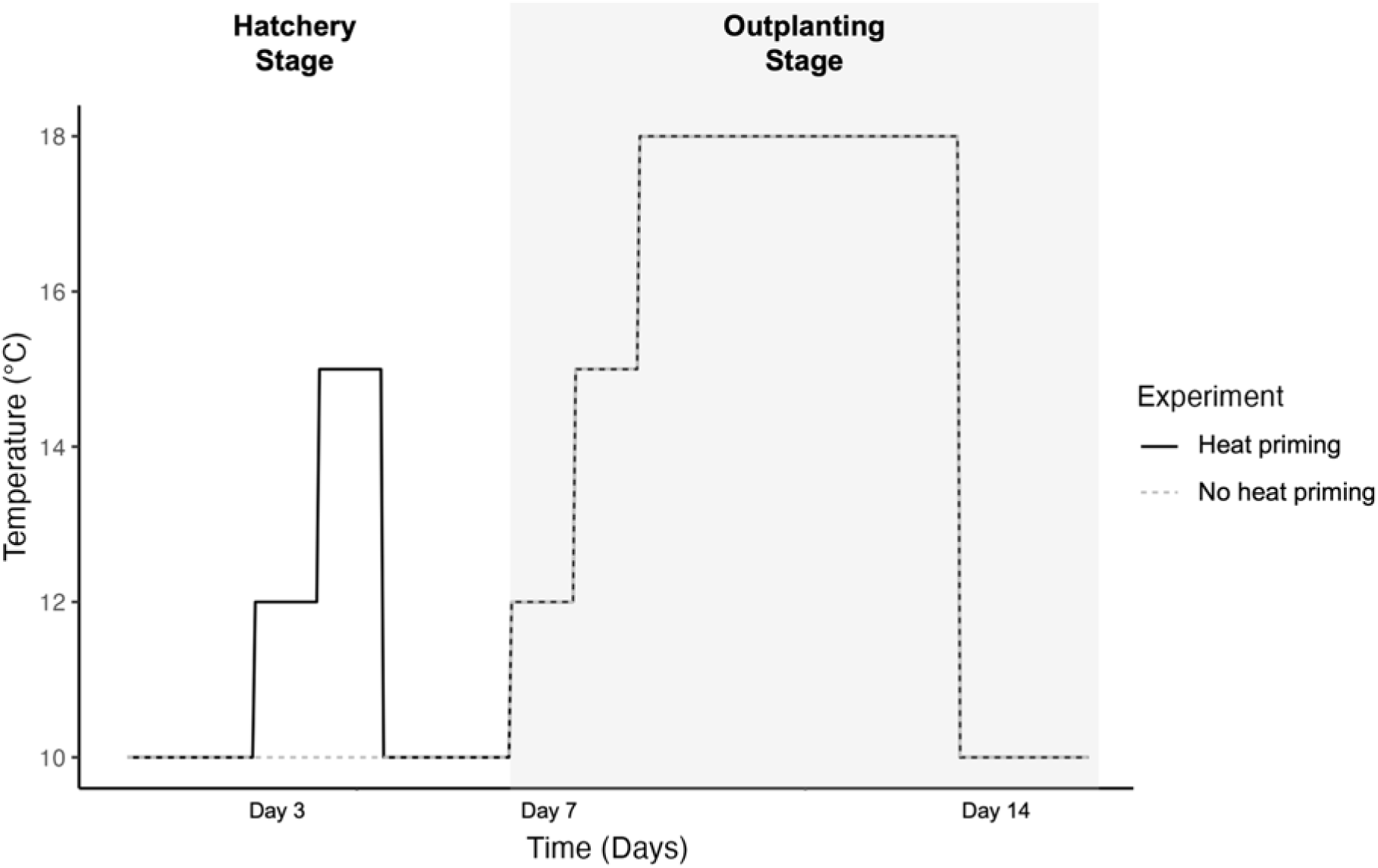
Temperature differences between heat priming treatments over the hatchery and outplanting stages Temperatures over the experimental window between heat primed (*solid line*) and non-heat primed (*dotted line*) treatments. During the hatchery stage, all seawater was UV treated and nutrients were added according to treatment assignment. During the outplanting stage, seawater was untreated, and no nutrients were added.

Again, the MHW level was informed by past thermal measures in the region of interest, specifically during the months outplanting typically occurs (November/December). The MHW was based on the common and minimum criteria of a defined heatwave event; at least five days in a row of temperatures exceeding 90% of the past observations at that location (46).

### 2.5 Kelp measurements

Wet weights and photographs of the spools were taken throughout the kelp performance and kelp priming experiments to determine differences in percent cover, coloration, and total weight across treatment and over time. Percent cover was taken to capture evenness of visible growth over the course of the experiment, while total weight of the spool allowed us to calculate relative growth rate. These combined measures provide insights into overall performance (growth and survival). Changes in coloration, or loss of pigment, can signal blade response to environmental stress(ors) (47,48).

At the conclusion of the experiment, all spools were unwound, a subset of blades were counted, and all sporophytes were removed. At this time chlorophyll-*a* fluorescence (*Fv/Fm*) was measured and individual blade photographs taken for color analysis (performance experiment only). All sporophytes were dried, weighed, and later set for stable isotope analysis. Blade counts can be used to compare blade density and overall success of each spool. Differences in chlorophyll-*a* fluorescence can indicate stress impacts on photosystem II (PSII), while an indirect effect of high temperature in kelp is loss of pigmentation, or bleaching (47). Blade measurements, such as surface area, show differences in growth and development between treatments. Dry weights, or dry content matter, indicate the total amount of growth on each spool. Any difference in measurements between the performance and priming experiment are noted below.

#### 2.5.1 Percent cover

To determine sporophyte percent cover of the mini-spools during both experiments, photographs (on the same side) were taken upon receipt of the spools (day 0), at the conclusion of the hatchery phase (day 7), and at the conclusion of the respective experiments (day 17). All photos were uploaded to and analyzed in Fiji Is Just ImageJ (FIJI) (49). Images were cropped to the growth area, converted to 8-bit in the Image menu and made binary in the Process menu (Fig 4). Black and white images were examined by an expert to ensure glares and shadows were removed, minor alterations were made when necessary, using the black and white paint tool. Finally, the Histogram tool from the Analyze menu was used to determine the number of black and white pixels (List function within Histogram), 0 indicating black and 255 white.

**Fig 4.**
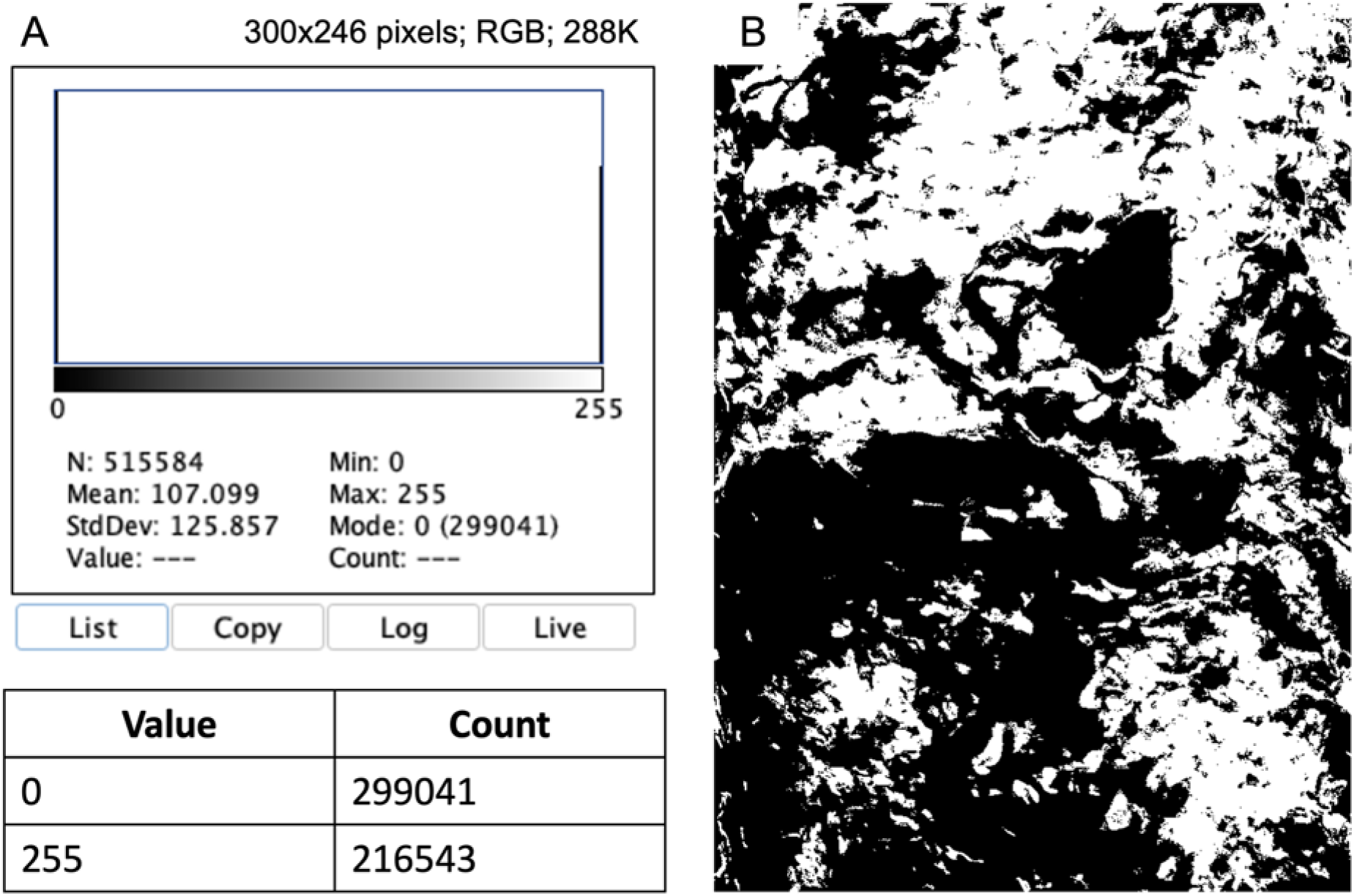
Percent cover measurement example from FIJI Example of percent cover measurement taken in FIJI for a mini-spool grown at 5°C during the outplanting phase of the 17-day kelp performance experiment. (A) Histogram of the color intensity within the mini-spool growth area with a list of numbers of black pixels (value = 0) and white pixels (value = 255). (B) A cropped black and white image of the growth area used to generate histogram in (A).

#### 2.5.2 *Weight*

We calculated the relative growth rates (RGR) of the mini-spools using a logarithmic formula for weight:

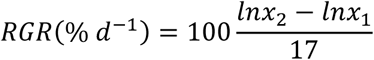

Spools were subsequently unwound and all sporophytic material collected, rinsed in fresh water, and dried to a consistent dry weight (g) at 60°C for at least 24hrs in a VWR drying oven for dry weight measures and isotopic analysis (see below).

#### 2.5.3 Blade count

To compare the density of blades between treatments at the end of the experiment, the unspooled twine was randomly cut into two 15cm sections from the middle of the spool. Each 15cm section was photographed and the blades were enumerated using the count tool in FIJI. There is no prescribed optimum density in current *M. pyrifera* aquaculture reports, thus our measurements potentially provide a comparative baseline.

#### 2.5.4 Chlorophyll-a fluorescence

We used a junior-PAM (Walz) chlorophyll-*a* fluorometer to measure the *Fv/Fm* of the juvenile sporophytes. *Fv/Fm* – a ratio that indicates the efficiency of photosynthesis based on maximum quantum yield of PSII and is commonly used as an indicator of plant stress – is the variable fluorescence divided by the maximum fluorescence. After at least 15 minutes of dark acclimation, *Fv/Fm* measurements were taken in the middle of the blade across ∼10 blades for each spool. We calculated the average of these *Fv/Fm* readings, resulting in a single value per spool.

#### 2.5.5 Coloration in performance curve

Given the importance of photosynthesis to the success of kelp growth and survival but limited capacity to measure in actual farm application, we set out to assess the relationship between coloration and photosynthetic efficiency across temperature treatments; an easier metric to potentially collect and assess (i.e., a photo). Kelps and other seaweeds are known to bleach when under stress (47), which can indicate a negative impact to photosynthesis (50). Measurements of mini-spool coloration were performed using FIJI on photographs at the end of the hatchery stage (day 7) and the end of the experiment (day 17) for the kelp performance experiment. The area of growth was outlined manually with the polygon tool and the histogram tool was used to determine the mean value for spool coloration (51) (Fig 5). Smaller values of coloration indicate darker pixels in the image.

**Fig 5.**
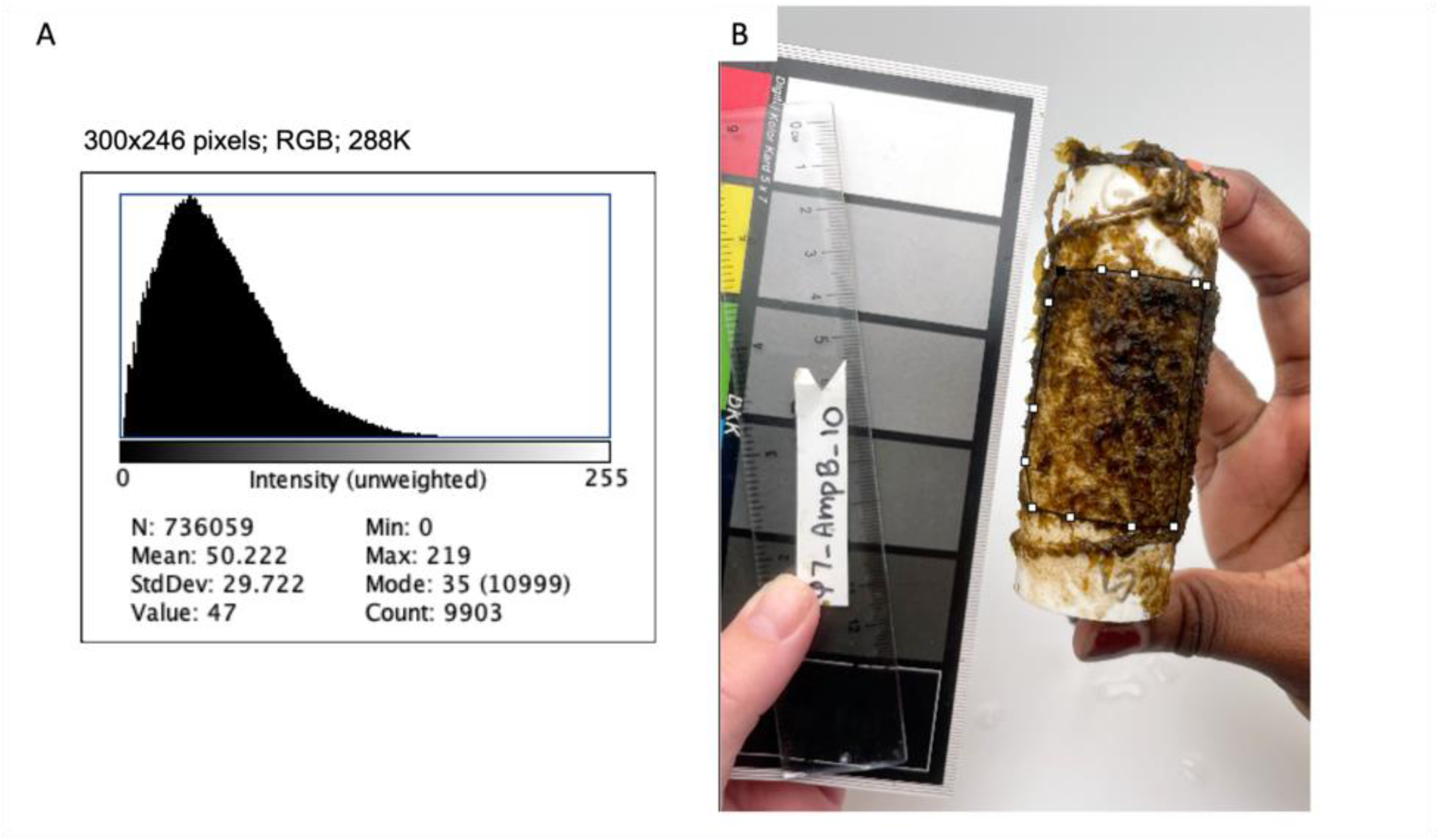
Coloration measurement example from FIJI Example of coloration measurement taken in FIJI for a mini-spool grown at 10°C during the outplanting phase, photo taken at the conclusion of the 17-day kelp performance experiment. (A) Histogram of the color intensity within the mini-spool growth area. (B) Outline of the growth area inside the polygon used to generate the histogram in (A).

#### 2.5.6 Blade surface area, length, and width in performance curve

For the kelp performance curve experiment we collected blade surface area, length, and width in FIJI. These measurements provide individual, trait-based assessments of growth to compare to the overall performance of the spool. When possible, five blades per spool were collected from the string tails (these tails were discarded and not used in the dry weight, coloration, or percent cover analysis). Blades were placed on a micrometer and images were taken using a dissecting scope. In FIJI, the scale was set in the Analyze menu using the micrometer. Surface area was calculated using the polygon tool to manually outline the blade, while length and width were determined with the straight tool (Fig6). Blade surface area, length, and width were averaged across mini-spool (N = 24).

#### 2.5.7 Nutrient content and stable isotope values

Tissue samples from each spool were collected and dried for stable isotope analysis. All samples were ground, weighted to the nearest microgram, and packaged in aluminum foil following the protocols for the UC Davis Stable Isotope Facility, where the analyses were performed (52). These results help inform how thermal stress may be impacting nutrient uptake and assimilation. More specifically, isotopic composition can reflect shifts in carbon fixation pathways, metabolic rates, nutrient use, and internal allocation of resources under different environmental conditions because biochemical reactions discriminate between isotopes in predictable ways (i.e., fractionation) (42,53). For instance, changes in δ¹³C in kelp tissue can indicate changes in photosynthetic carbon assimilation or shifts in the source of carbon (54,55). Our samples were analyzed for total carbon content (%), total nitrogen content (%), δ^13^C, and δ^15^N using an EA-IRMS system at the UC Davis Stable Isotope Facility.

### 2.6 Statistical analysis

#### 2.6.1 Kelp performance curve

To analyze the reaction norms of the proportion of mini-spool coverage, relative growth rate, final dry weight, blade count and *Fv/Fm* across the temperature gradient (day 17), we compared several generalized linear models – specifically an intercept, linear, and second order polynomial regression – using a beta distribution with a logit link function and calculated the associated Akaike Information Criterion Corrected (AICc) values. The beta distribution was selected because proportion is bounded between 0 and 1. AICc allows a likelihood comparison of model fit in relation to model complexity (56,57). Best fit was determined by the model with the lowest AICc value. The R package *glmmTMB* was used to build the comparative models, while the *aictab* function in the package *AICcmodagv* was used to compare model performance (58,59). We further tested the effect of time (via day 0, 7, 17) across the gradient on coverage to discern when significant changes were detectable, if any.

To assess how color related to photosynthetic performance, we visualized the difference in color intensity between day 7 to day 17 relative to the corresponding *Fv/Fm*. We followed this with a Kruskal Wallis test to compare *Fv/Fm* levels between replicates that clustered together.

To discern how temperature impacts kelp growth on an individual level, we performed one-way analysis of variance (ANOVA) for blade surface area, height, and width as residuals were normal and variances homogeneous. We followed those with Tukey’s post hocs.

We also assessed the relationship between % C by % N, fit with a linear model. Such a comparison provides insights into the range and relationship for these earlier life stages. This was followed by a post hoc cluster analysis to determine if the relationship between carbon and nitrogen grouped by temperature exposure treatment.

Percent carbon, carbon to nitrogen ration, δ^13^C, and δ^15^N met normality assumptions for parametric statistics and measures across the six treatments were analyzed using one-factor ANOVAs followed by Tukey’s post-hoc test. A Kruskal-Wallis followed by a Dunn’s post hoc was used to analyze %N due to normality assumptions not being met.

#### 2.6.2 Priming giant kelp

In addition to the differences between phase 1 and phase 2 described above, there was a bacterial outbreak during phase 2 before the spools were transported to UC Santa Barbara. Due to this outbreak, water was changed more frequently at Ocean Rainforest and the sporophytes were a few days older than phase 1 sporophytes at the start of the experiment. ‘Phase’ was too even to be added as a random effect in our models, so instead we compared the two phases with appropriate post-hoc analyses.

To assess the comparative impact of our four treatments [controls (unprimed), heat primed, nutrient primed, and fully (heat + nutrient) primed] on the proportion of mini-spool coverage over time (day 0, 7, and 17), we followed the same steps described for the performance curves. We also conducted a two-factor ANOVA on day 0 proportions to ensure no significant differences at the start of the experiment, given the phased approach we had to employ due to tank-space limitations.

The other ‘end of experiment’ metrics (day 17) – blade count, relative growth rates, *Fv/Fm*, and dry weight – were visualized and regressed with appropriate distributions relative to the data type. Since time was not a factor, the various models only compared the differences of the four treatments. Final dry weight met normality assumptions and was analyzed using a linear model. For the other metrics, the *glmmTMB* package was used to assess the significance of any differences. For blade count, we used a gamma distribution with a log link function. Relative growth rates were compared using a gaussian distribution and an identity link function, while we used a beta distribution with a logit link function for *Fv/Fm.* All models were followed by post-hoc tests calculating the least-squares means using the function *emmeans* to determine pair-wise differences between treatments.

Percent carbon, percent nitrogen, and carbon to nitrogen ratio met normality assumptions for parametric statistics and primed treatment effects were analyzed using linear models. To assess differences in δ^13^C we used a glmmTMB with a gaussian distribution and an identity link function. For all nutrient analyses, we once again conducted appropriate post-hoc tests to compare phase 1 and phase 2 trials. Due to obvious phase differences, for δ^15^N we only conducted Kruskal Wallis comparing phase 1 and phase 2.

All statistical analyses across the two experiments were performed in R version 4.2.3 (60).

## RESULTS

At the conclusion of the kelp performance experiment, 22 HOBO pendants successfully logged temperatures across the 17 days. Most experimental tanks had two replicate loggers, however two loggers from different tanks failed to record. The average temperature across all tanks before the heatwave treatment began was 10.17°C (± 0.002). During the 7-day heat treatment, average temperatures ranged from 7.21°C (± 0.012) for the 5°C treatment to 28.40°C (± 0.042) for the 30°C treatment (Table 1 and S1 Fig). After the heat treatment, the average temperature across all experimental tanks was 10.24°C (±0.006) for the 2-day recovery period.

**Table 1.** Temperatures across kelp performance experiment.

| Treatment | Actual Temperature |
| --- | --- |
| $5^\circ\text{C}$ | $7.21^\circ\text{C}$ ($\pm 0.0115$) |
| $10^\circ\text{C}$ | $11.22^\circ\text{C}$ ($\pm 0.0050$) |
| $15^\circ\text{C}$ | $16.03^\circ\text{C}$ ($\pm 0.0103$) |
| $20^\circ\text{C}$ | $20.02^\circ\text{C}$ ($\pm 0.0246$) |
| $25^\circ\text{C}$ | $24.55^\circ\text{C}$ ($\pm 0.362$) |
| $30^\circ\text{C}$ | $28.40^\circ\text{C}$ ($\pm 0.0424$) |

At the conclusion of phase 1 and 2 of the kelp priming experiment, 21 out of 24 HOBO pendants successfully logged across the 17 days. Temperatures remained close to the target temperatures for each phase (Table 2 and S2 & S3 Figs).

**Table 2.** Actual temperatures for each treatment in the priming experiment.

| Stage |  | Not Primed | Heat Primed | Nutrient Primed | Fully Primed |
| --- | --- | --- | --- | --- | --- |
| Hatchery | Acclimation | $10.27^\circ\text{C}$ ($\pm 0.0015$) | Did not record | $10.09^\circ\text{C}$ ($\pm 0.0030$) | Did not record |
| | Heat prime day 1 | | $13.04^\circ\text{C}$ ($\pm 0.0129$) | | $13.08^\circ\text{C}$ ($\pm 0.0124$) |
| | Heat prime day 2 | | $15.63^\circ\text{C}$ ($\pm 0.0170$) | | $15.84^\circ\text{C}$ ($\pm 0.0111$) |
| | Recovery | | $10.18^\circ\text{C}$ ($\pm 0.0208$) | | $10.45^\circ\text{C}$ ($\pm 0.0185$) |
| Oceanic | Trigger day 1 | 13.11°C(±0.0108) | 12.98°C(±0.0116) | 12.98°C(±0.0116) | 13.11°C(±0.0108) |
|  | Trigger day 2 | 15.84°C(±0.0085) | 15.61°C(±0.0130) | 15.61°C(±0.0130) | 15.84°C(±0.0085) |
|  | Trigger days 3-7 | 18.45°C(±0.0025) | 18.28°C(±0.0042) | 18.28°C(±0.0042) | 18.45°C(±0.0025) |
|  | Recovery | 10.47°C(±0.0128) | 10.39°C(±0.0150) | 10.39°C(±0.0150) | 10.47°C(±0.0128) |

The actual temperatures (±SE) for each treatment in the priming experiment across the Hatchery and Oceanic stages. Phase 1 includes the ‘not primed’ and ‘fully primed’ treatments, while phase 2 includes the ‘heat primed’ and ‘nutrient primed’ treatments.

### 1.1 Kelp performance curve

Across all performance measures we found optimum patterns typically between 10-15°C and a clear threshold response when temperatures exceeded 20°C (Fig 7). In fact, for spool coverage, relative weight, and final dry weight we did not include the 25 and 30°C temperature treatments due to cyanobacterial growth but did include them for average blade count where we could indicate zero blades and average *Fv/Fm* where we took readings from the few blades that survived. The best fit for the proportion of mini-spool coverage on the final day was a polynomial regression (AICc = -17.01 , AICcWt = 0.57; R^2^ = 0.31, F-stat = 24.41 on 2 and 21 DF), peaking at 15°C at 0.79 (±0.06) cover (Fig 7A). For relative growth rate, the best fit was also a polynomial regression (AICc = -33.30, AICcWt = 0.79 ;R^2^ = 0.41, F-stat = 6.6136 on 2 and 13 DF) with the peak occurring closer to 10°C, which had an average of 30.2% d^-1^ (±0.02) (Fig 7B). The best fit for final dry weight was actually a linear model (AICc = -56.74, AICcWt = 0.64; R^2^ = 0.10, F-stat = 2.608 on 1 and 14 DF). The dry weights where highest for the 5°C and 10°C treatments (0.12g ± 0.01 and 0.15g ± 0.02, respectively), then decreased by an average 0.02g (±0.01) per ca. 5°C as temperatures increased (Fig 7C).

**Fig 6.**
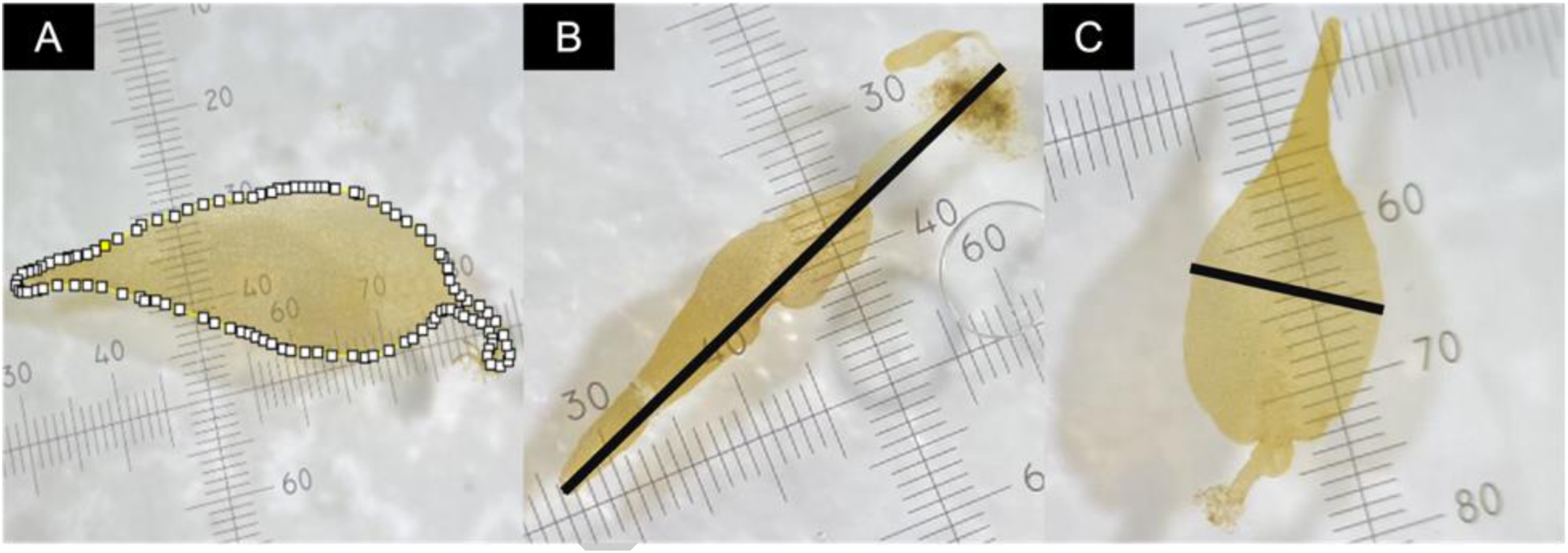
Examples of blade measurements taken in FIJI. (A) Surface area of blade. (B) Length of blade. (C) Width of blade.

**Fig 7.**
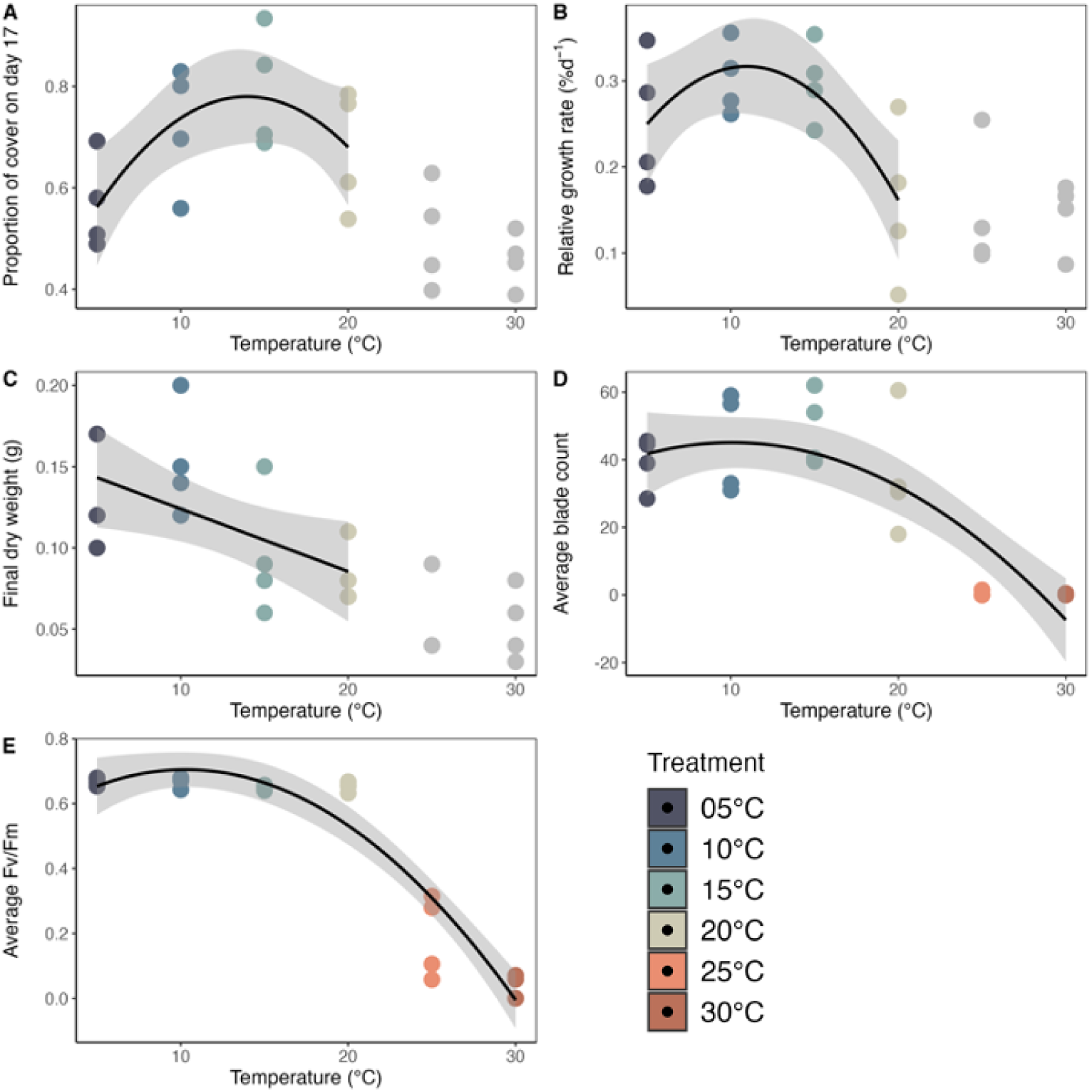
Kelp performance experiment outcomes (A) The proportion of mini-spool coverage on the final day of the experiment fit with a polynomial regression. (B) The relative growth rate (%d^-1^) fit with a polynomial regression. (C) The final dry weight (g) fit with a linear model. (D) The average blade count per 15cm fit with a polynomial regression. (E) Average *Fv/Fm* fit with a polynomial regression. For all panels, the black line represents the best fit model surrounded by grey 95% confidence intervals. Grey points in panel A-C are shown for reference but not included in the model due to overgrowth of cyanobacteria, thus inability to distinguish between kelp versus bacterial biomass.

Patterns at the spool (population) level mimicked those at the individual level. For average blade count, the best model fit was the polynomial regression (AICc = 198.47 , AICcWt = 0.95; R^2^ = 0.67, F-stat = 4.369 on 2 and 21 DF). The 15°C treatment had the highest blade count, with an average of 49 blades per 15cm, however the counts were similar for the 10°C treatment (Fig 7D). The best fit for *Fv/Fm* was a polynomial regression (AICc = -28.82, AICcWt = ; R^2^ = 0.88, F-stat = 87.7 on 2 and 21 DF) (Fig 7E). Any bleaching was noted qualitatively but was avoided when determining the final *Fv/*Fm values, as we were interested in the intact photosynthetic tissues. Several bleached spots were tested, resulting in a zero every time. This does not inform the overall photosynthetic potential of a mini-spool. The average *Fv/Fm* for the cooler temperature treatments (5 – 20°C) ranged from 0.649 – 0.669, with a precipitous drop off for the higher temperature treatments.

Modeling the full extent of the proportional coverage, the best fit model, based on the AICc value and weight, (AICc = -116.21, AICcWt = 0.78; see S4 Fig, S1 & S2 Table) included day and treatment and the interaction between the two. There was specifically a significant interaction between the final day (day 17) and two temperature treatments,10°C and 15°C (p-values = 0.007 and < 0.001, respectively), indication that coverage evenness was significantly higher between these two thermal conditions after recovery (day 17), but not before.

There was a clear threshold delineation between the highest temperature treatments (25°C and above), including coloration (Fig 8). Across the 5 – 20°C treatments, the majority (88%) of mini-spools decreased in color intensity (i.e., became darker), whereas all warmer treatments saw an increase (Fig 8A). The cooler treatments (≤ 20°C) formed a tight aggregation between 0.640 to 0.680 *Fv/Fm* and displayed a difference in color intensity less than 22 (Fig 8B). In comparison, the warmer treatments had readings below 0.400 and above the 22-coloration difference threshold. A post-hoc analysis comparing *Fv/Fm* of these two clusters found significant differences (Kruskal-Wallis; p-value < 0.001), with *Fv/Fm* decreasing by an average of 0.370.

**Fig 8.**
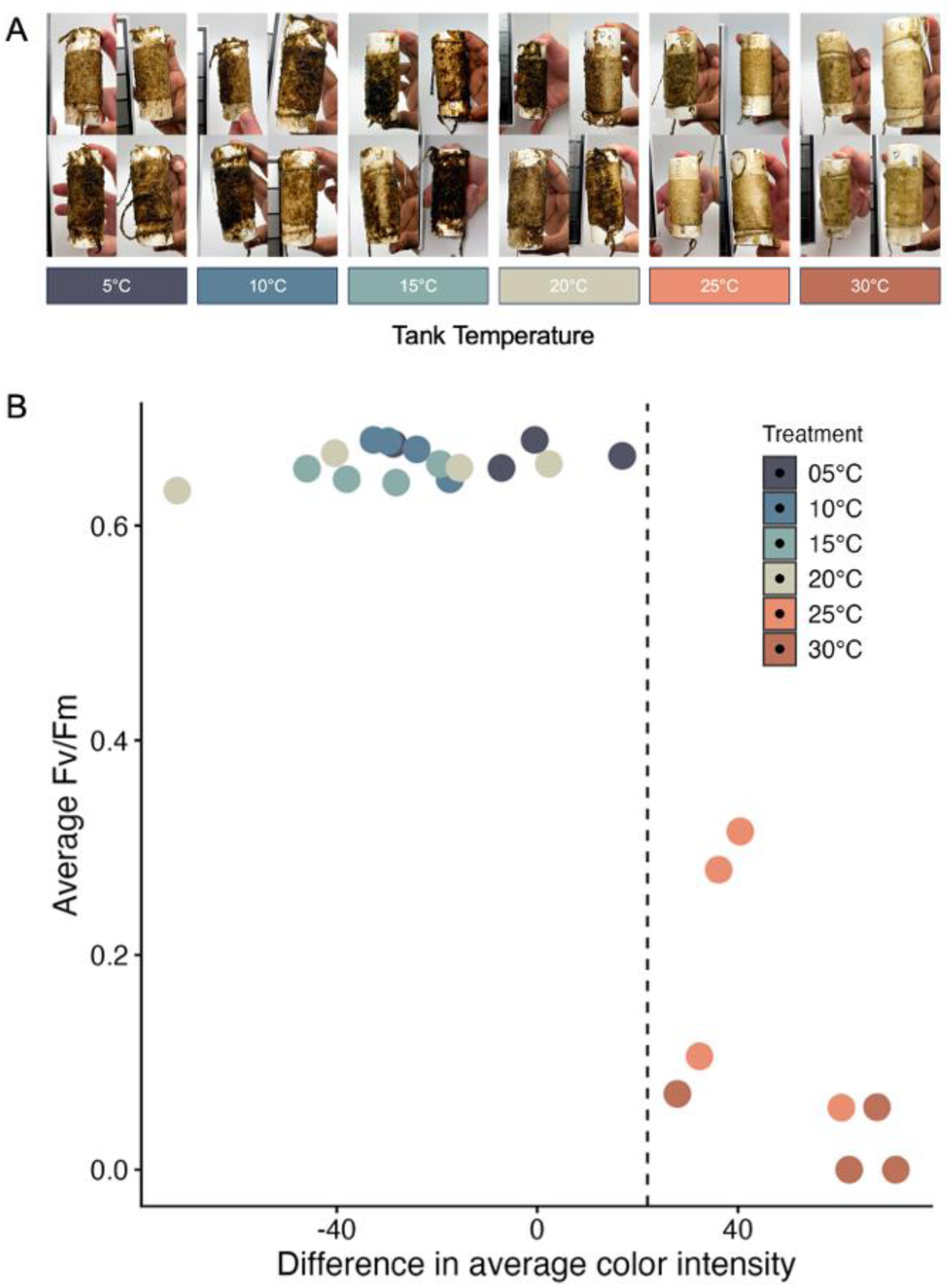
Average *Fv/Fm* plotted against the difference in average color intensity. (A) A photo of each mini-spool organized by their treatment temperature and visual representation of the change in coloration relative to temperature. (B) Quantitative relationship of the mean *Fv/Fm* and mean difference in color intensity between day 7 and day 17 [positive values = lighter (bleaching); negative values = darker]. The dotted line (color intensity = 22) indicates the threshold color associated with a significant reduction in photosynthetic efficiency above 20°C.

Overall, blade growth across all metrics was restricted at temperatures at or above 20°C (Fig 9), similar to the broader mini-spool measurements trends. Average surface area was highest at 15°C and lowest at 25°C (Fig 9A; one-way ANOVA, p-value = 0.0137). Average height decreased at higher temperatures starting at 20°C (Fig 9B; one-way ANOVA, p-value = 0.0012), while average width decreased at temperatures above and below 15°C (Fig 9C, one-way ANOVA, p-value = 8.72x10^-5^). We also found fewer blades at higher temperatures, with only three blades found total on the 25°C mini-spool tails and no blades in the 30°C treatments (i.e., 100% mortality).

**Fig 9.**
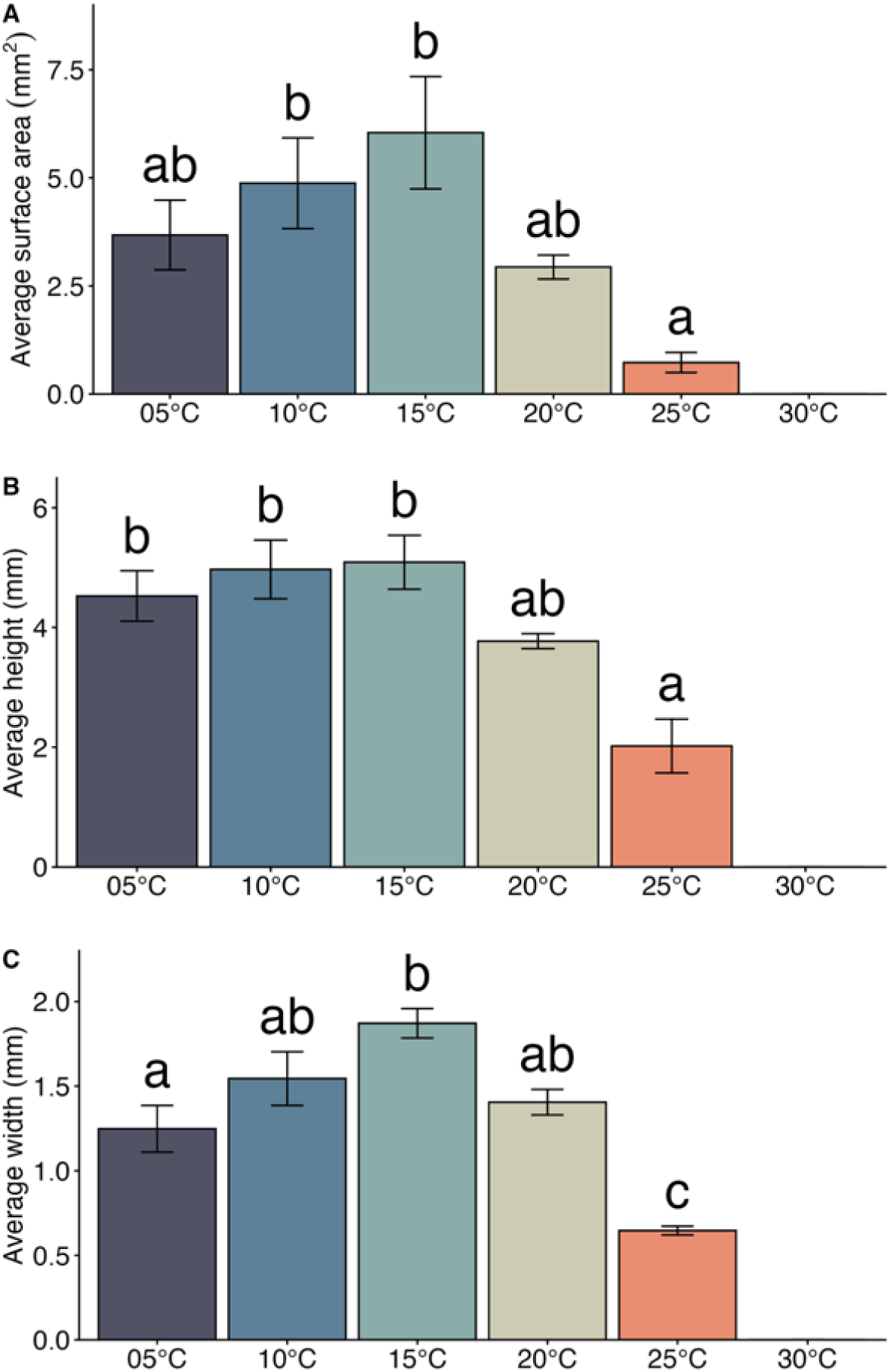
The impact of different temperature regimes on blade growth. (A) Average surface area, (B) average height, and (C) average width across the temperature treatments. Bars are means ±SE. Bars with different letters are statistically different.

There were not significant differences between treatments for percent carbon (one-factor ANOVA, p-value = 0.0964; Fig 10A) or percent nitrogen (Kruskal-Wallis, p-value = 0.06017; Fig 10B), but there were differences in the carbon to nitrogen ratio (one-way ANOVA, p-value = 0.0016, Fig 10C).

**Fig 10.**
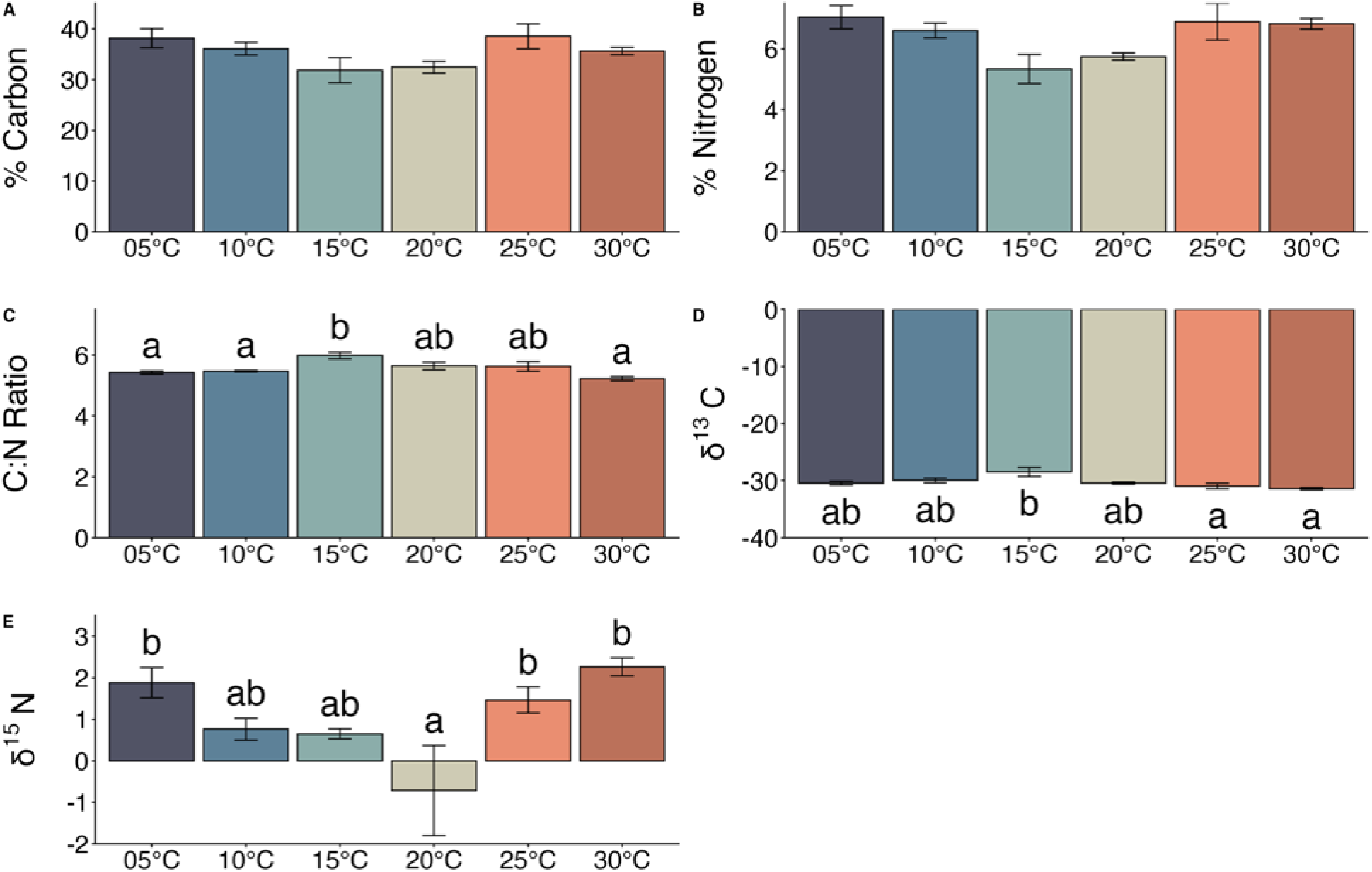
Nutrient analysis across temperature regimes. (A) Percent carbon, (B) percent nitrogen, (C) carbon to nitrogen ratio, (D) δ^13^C, and (D) δ^15^N across temperature treatments. Bars are means ±SE. Bars with different letters are statistically different.

Reflective of the broader trends, the C:N ratio was the highest at the 15°C treatment and the lowest at 5, 10, and 30°C. The average ratio, however, at 15°C was only 0.76 parts carbon to nitrogen larger than 30°C treatment (Fig 10C). There were also significant differences in δ^13^C (one-way ANOVA, p-value = 0.0072) where the 15°C treatment is higher than the 25 and 30°C treatments (Fig 10D). δ^15^N had significant differences across treatments (one-way ANOVA, p-values = 0.0032) with the 20°C treatment lower than the 5, 25 and 30°C treatments (Fig 10E).

As expected, percent nitrogen increased linearly with percent carbon (R^2^ = 0.91, F-stat = 202.8 on 1 and 20 DF, Fig 11) with values ranging from 4.14 to 8.02% for N and 26.11 to 43.75% for C. We performed a post hoc cluster analysis to examine patterns in this relationship across temperatures, however extreme heat did not stand out (see S5 Fig). For all nutrient and isotope analyses, it is important to note two samples were not included (one from both 5°C and 20°C treatments) as the samples were too small for isotopic analysis.

**Fig 11.**
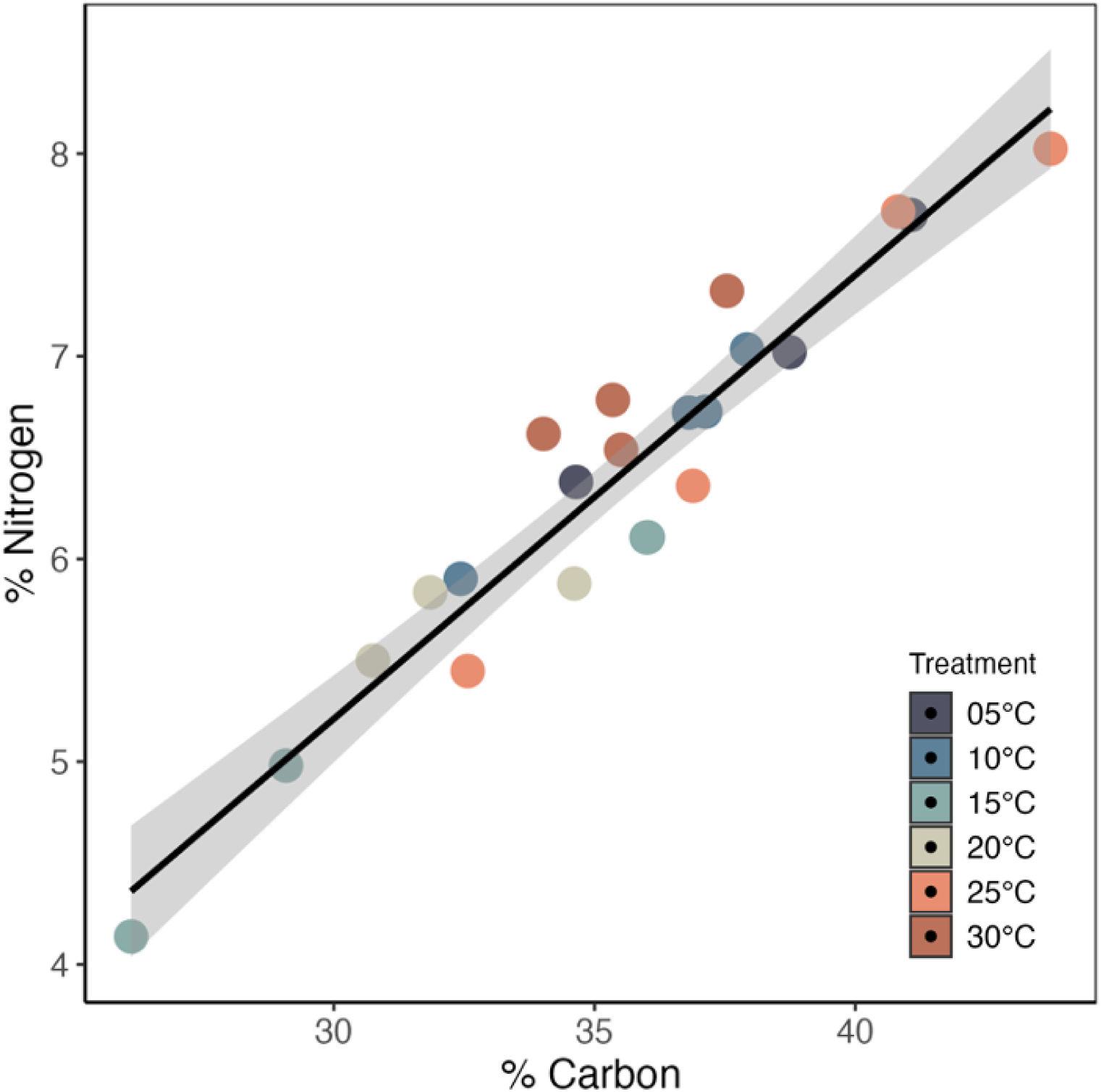
Relationship between percent carbon and percent nitrogen. There is a linear relationship between percent carbon and percent nitrogen where the line represents the linear model and is surrounded by grey confidence intervals.

### 1.2 Priming giant kelp

The best fit model to examine differences in mini-spool cover over time included day, nutrient treatment, and heat treatment as fixed factors and their interactions (AICc = -242.8, AICcWt = 0.54; see S3 Table), with a significant three-way interaction between day, nutrient treatment, and heat treatment (p-value < 0.001, Fig 12 & S4 Table). On the final day, fully primed mini-spools had higher coverage than expected from the additive effects of nutrient and heat priming. There were no significant differences in proportion of cover at the start of the experiment (two-factor ANOVA, p-value = 0.4238) or between phase 1 and phase 2 (T-test, p-value = 0.9279).

**Fig 12.**
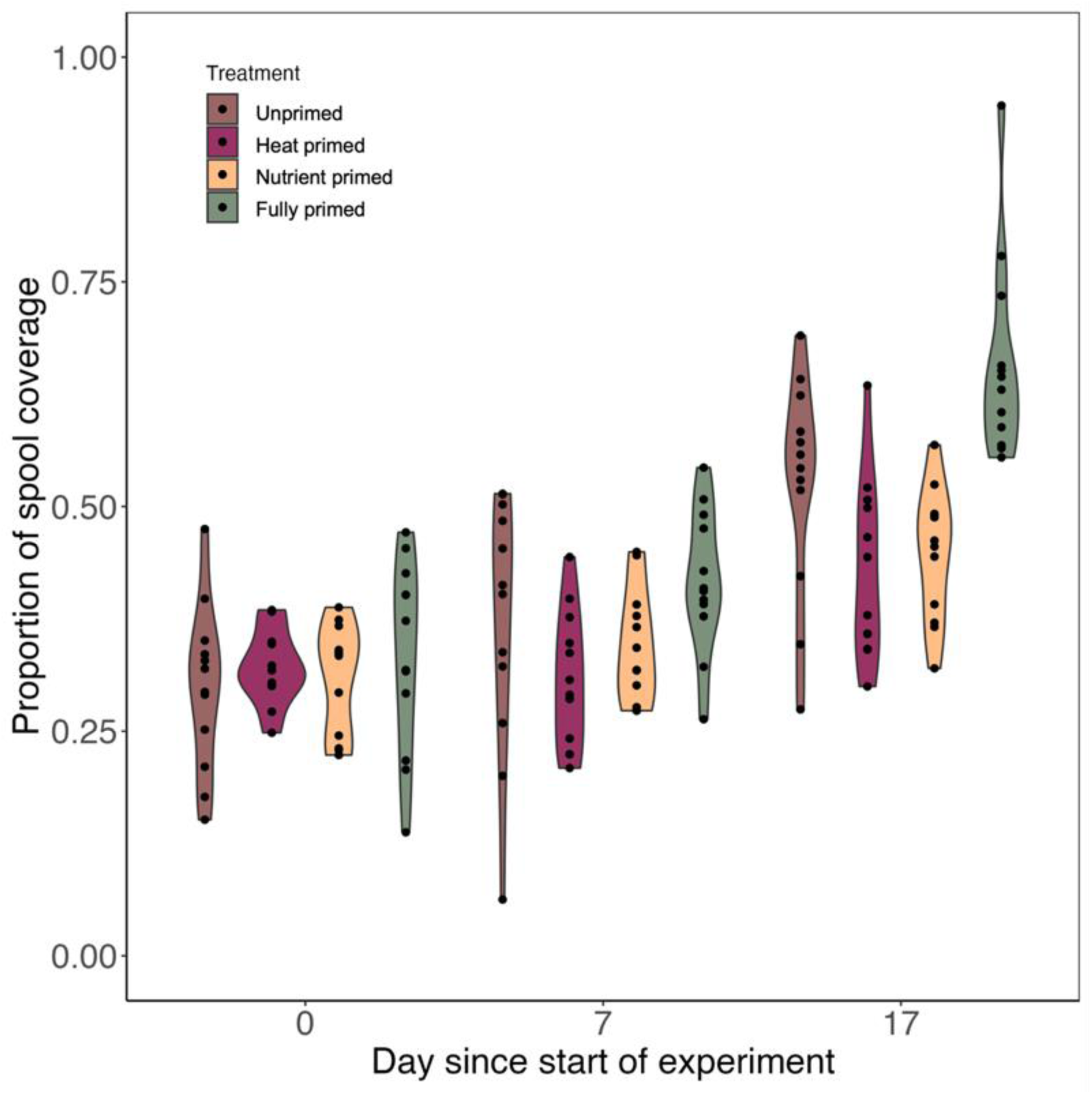
Change in coverage proportion over time for the priming experiment. Proportion of mini-spool coverage at the start of the kelp priming experiment (Day 0), after the hatchery stage (Day 7), and at the conclusion of the experiment (Day 17).

Fully priming (nutrients and heat) kelp sporophytes resulted in higher growth rates than single priming treatments (Fig 13A), and all priming led to higher dry weights than the control (Fig 13B). All factors were significant, and there was an interaction between factors when comparing relative growth rate (glm; p-value = 0.0075), with a significantly higher rate of growth for the fully primed mini-spools (mean = 0.478 %d^-1^) compared to only heat (mean = 0.266%d^-1^) or nutrient primed mini-spools (average = 0.303%d^-1^; Fig 13A). There were significant differences across treatments for final dry weight (linear model; p-value < 0.001). The unprimed treatment (control) had the lowest final dry weight out of all treatments, with the fully primed and nutrient primed treatments weighing over 2x more (Fig 13B). All factors were significant for average blade count per 15cm (glm; interactive p-value < 0.001) with the unprimed and fully primed treatments (phase 1) having more blades per 15cm (Fig 13C). The average *Fv/Fm* also had a significant interaction between the factors (p-value < 0.001), with the fully primed and unprimed treatments (phase 1) having ∼8 times higher blade counts (Fig13D). Post-hoc analyses found an effect of phase for each metric.

**Fig 13.**
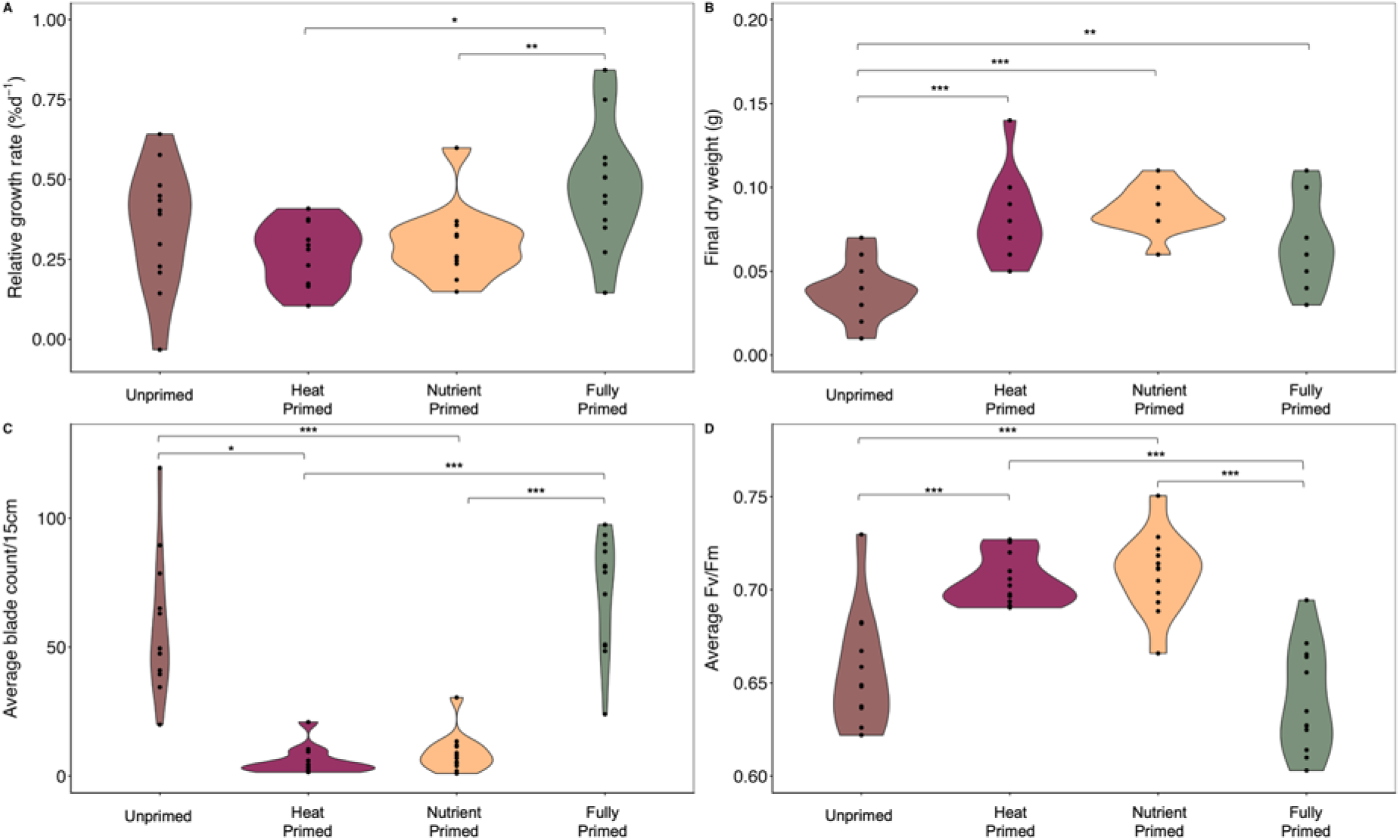
Kelp priming experiment outcomes The difference across treatments for (A) the relative growth rate (%d^-1^), (B) final dry weight (g), (C) average blade count per 15cm, and (D) average *Fv/Fm*. The bars show significant differences between treatments where * < 0.5, ** < 0.01, and *** < 0.001.

There were not significant differences between treatments for percent carbon (linear model, p-value = 0.1698), percent nitrogen (linear model, p-value = 0.1063), or carbon to nitrogen ratio (linear model, p-value = 0.405; Fig14A – C). Post-hoc analyses found a trial effect for the carbon to nitrogen ratio; however, all values were between 5.9 and 7.4 (difference of 1.5). All factors were significant and there was a significant interaction between factors for δ^13^C (glm, interactive p-value < 0.001). The heat primed and nutrient primed (phase 2) had higher values than the other treatments (Fig 14D). Post-hoc analysis confirmed a phase effect for δ^13^C (Kruskal Wallis, p-value < 0.001). δ^15^N values were positive for the phase 2 treatments and negative for the phase 1 treatments (Fig 14E). A Kruskal Wallis comparing phase 1 with phase 2 found significant differences (p-value < 0.001). For all nutrient and isotope analyses, it is important to note three samples were not included (three from the control treatment) as the samples were too small for isotopic analysis.

**Fig 14.**
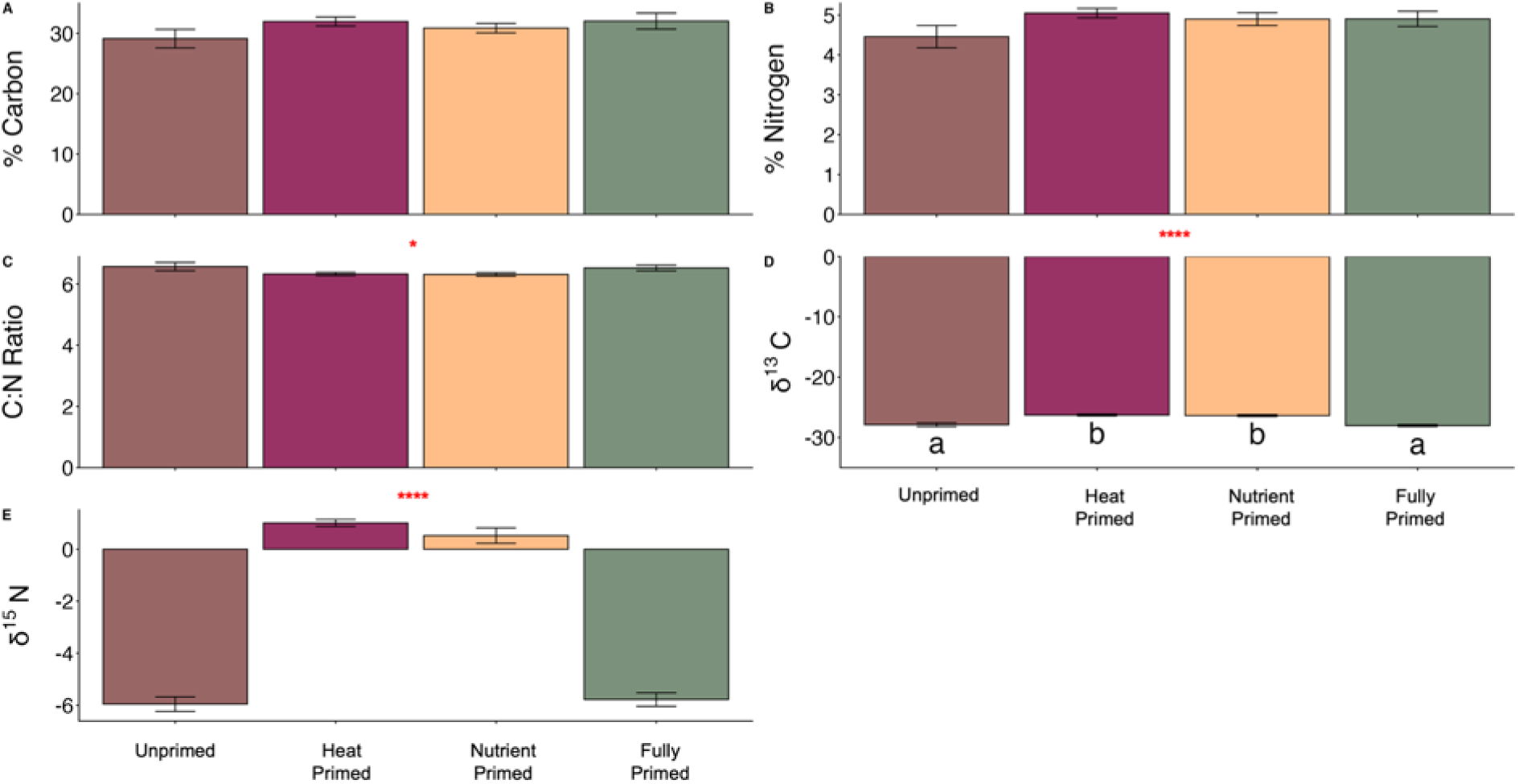
Nutrient analysis across priming treatments. (A) Percent carbon, (B) percent nitrogen, (C) carbon to nitrogen ratio, (D) δ^13^C and (D) δ^15^N across priming treatments. Bars are means ±SE. Bars with different letters are statistically different. Red asterisks indicate a phase effect was found in post hoc analyses where * < 0.5, ** < 0.01, *** < 0.001, and **** < 0.0001.

## DISCUSSION

We make several important contributions toward understanding the temperature tolerance and capacity for priming in *M. pyrifera* at the juvenile sporophyte stage. First, fully priming this species in a hatchery setting resulted in overall increased performance during a MHW, lending support that priming can be a viable intervention for aquaculture practitioners growing this species. Second, temperatures above 20°C had a significant negative impact on sporophytes during this early stage. Furthermore, our growth, survival, and physiological findings generally support the current outplanting strategy in California of waiting for temperatures to reach 15°C but suggest cooler conditions (closer to 10°C) would be more favorable. These findings broaden our understanding of this species and support further research on priming seaweeds for climate stressors.

Fully priming (nutrients and heat combined) *M. pyrifera* mini-spools during the hatchery stage resulted in significantly higher coverage and relative growth after the simulated 18°C MHW, providing the first supportive evidence that priming could be a climate tool for aquaculture and restoration of the species. Although a different life stage and species, our results are similar to Gauci et al. 2024 (41), where authors found that *Saccharina latissima* (sugar kelp) sporophytes, another brown kelp common in aquaculture, grew 30% faster when gametophytes were primed for 4 weeks before spores were created.

Gauci et al. 2024 (41)’s results lend support to potentially longer lasting priming, possibly through epigenic DNA-methylation; though this was not explicitly tested. Our priming was considerably shorter – days instead of weeks – which is more common in the literature (19). We also focused on juvenile sporophytes given the dearth of information on this critical transitional stage and the uncertainty of developmental reset. Even so, we still found comparable growth (average of 0.35% day^-1^ across treatments) to Gauci et al. 2024 (41), almost a 15% increase in spool coverage for fully primed spools, and better evenness across the spools, indicating it was not a function of just selection of the best growers. Gauci et al. 2024 (41) did not, however, test nutrients, which we found was likely important in improving tolerance of our species, similar to Fernández et al. 2020 (61) and in support of the *nutrient rescue hypothesis* in *M. pyrifera*. The nutrient rescue hypothesis states high nutrient availability can alleviate impact from stressors, allowing individual to compensate for the energetic cost of the stress (62,63).

For both the kelp performance and priming experiments, the nitrogen levels (5.3% ±0.18) were well above reported values in other studies of *M. pyrifera* (39,64), a function of a farmed versus wild setting. Bunting et al. 2024 (42) reported nitrogen averaged below 1.8% across all treatments. Our experiment surpassed nitrogen levels above the saturation point found by Wheeler and North 1980 (65), where external nitrite exceeded 25μg of N liter^-1^. This was true across both experiments, which each had one week with hatchery level nutrients or half-hatchery level nutrients and no added nutrients during the outplanting phase. The increased level of nitrogen may have contributed to overall improved thermal performance of this species, as other research on *M. pyrifera* adult sporophytes also found nutrients to enhance growth and photosynthesis at higher temperatures (39). This suggests that the industry standard has more than enough nitrite for *M. pyrifera* hatcheries, with levels remaining elevated for at least 10 days post outplanting and even up to two weeks (66). However, further research should be conducted to determine which species of kelp experience nutrient rescue at higher temperatures, as increased tolerance does not appear to be ubiquitous. Elevated nutrients did not buffer the effects of high temperatures for gametophyte or early sporophyte stage *Nereocystis lutekeana* (bull kelp), a kelp that can be found from central California up into Alaska (67) and has seen significant declines caused by MHWs (68). Another study on the sporophyte blades of *N. lutekana* and *S. latissima* found added nitrogen did not improve growth at 21°C for either species, and in fact found increased biomass at lower temperature (13 and 16°C) when grown under “low” nitrogen conditions (69). Timing of nutrient addition might also play a role, as Fales et al. 2023 (69) also found that blades grew more biomass across temperatures when sourced from a site with previously higher dissolved inorganic nutrients. This suggests a possible lag effect and future studies should focus on the timing of nutrient addition on temperature amelioration.

We found temperatures above 20°C had major negative impacts across all measures, affecting growth, survival, and physiology of juvenile *M. pyrifera* sporophytes at the individual and population (spool) level. Although a few blades were able to survive at temperatures above 20°C, these blades had reduced photosynthetic capacity, were smaller in size, and showed clear evidence of bleaching. Similar temperature thresholds have been reported at other life stages of *M. pyrifera*. For example, in New Zealand, gametophytes experience inhibited fertilization above 18.8°C and complete secession of fertilization at 23.6°C (70). Similarly, a study on populations in California and Chile found gametophytes survived at 20 and 18°C respectively but Chilean gametophytes produced no eggs, while Californian gametophytes produced some eggs, but no sporophytes developed (38). A study modeling future climate conditions predicts precipitous declines in the future at southern and northern California sites, with complete extirpation of *M. pyrifera* at the southern site by 2100 (71). Furthermore, the thermal optimum (based on growth and photosynthesis) for adult sporophytes range between 14 and 19°C, with CTmax of 26°C (39). In our study, however, even 20°C resulted in some metrics declining (coverage, growth rate, dry weight, surface area, height and width). These findings provide evidence that (unprimed) juvenile sporophyte sensitivity maybe more similar to gametophytes than their adult sporophyte counterparts.

Given this stage is the one that experiences real ocean temperatures first in aquaculture and restoration settings, this higher sensitivity will be important in considering the climate change consequences on the species in the future.

While our findings of the performance curves generally support the current practice for *M. pyrifera* in California aquaculture of outplanting in November or December at 15°C (Ocean Rainforest personal correspondence), the increasing occurrence and intensity of MHWs in the region highlights impending threats (72). We did find the proportion of mini-spool coverage, the relative growth rate, and the average blade count to be highest at 15°C; although some measures were equivalent or better closer to 10°C (e.g., RGR, final dry weight, *Fv/Fm*), suggesting cooler temperature are better for this stage. Notably, these were unprimed measures and we found metrics began to decline even under 20°C. Chan et al. 2024 (72) recently assessed spatial and temporal MHW patterns in the Santa Barbara channel, finding anomalous thermal conditions occur year around, including 2-3°C average above normal in the winter months in more years. In fact, the average near-surface conditions during November and December are trending above 15°C, reaching more sustained levels closer to 18°C during the 2015 blob. Long term, Fong et al. 2024 (35) projected a decrease in seaweed aquaculture performance under future climate conditions, however these projections do not account for climate extremes or changes in farming practices. Thus, as ocean temperatures continue to rise and heatwaves become more common, maintaining the current outplanting schedule will likely need to be reassessed. In the near future, outplaning may need to be delayed into January and/or other adaptive measures adopted, including priming. Genetic selection could theoretically help (73), but is currently prohibited in the West Coast of the U.S. and thus presently not a viable option unless policies change.

We found that blade color directly reflects the performance of photosynthesis in *M. pyrifera* and could potentially be used on kelp farms as an indicator of health. The measurable lightening of color intensity we observed in the higher temperature treatments corresponded to a threshold response with significantly lower *Fv/Fm* when difference in color intensity exceed 22. As it may be more expensive and time consuming for farmers to directly test photosynthesis, color grading could be a simple and non-invasive way for farmers to assess their crop over time via visual inspection. Color grading is extremely common in terrestrial agriculture, including for strawberries, tomatoes, apples, and soybeans, to name a few (74–76). In fact, Nori (*Pyropia spp.*) – seaweed commonly used in sushi for direct human consumption – has a color system in place (77). Color scores such as these are also used in assessing coral health, both by researchers and public science initiatives (78,79). Thus, a simple color chart could be developed from our work and used for a quick visual assessment tool, with the added benefit of knowing the relationship to photosynthetic performance.

Despite no significant differences in bulk percent C or N, the temperature-driven changes in C:N and stable isotopes in the performance experiment provide evidence that thermal stress alters carbon acquisition strategy rather than total nutrient content, and priming may be beneficial. The elevated C:N and higher δ¹³C at 15°C suggest more efficient inorganic carbon uptake consistent with active carbon-concentrating mechanisms (CCMs) under near-optimal conditions (80,81). In contrast, the more negative δ¹³C values at 25-30 °C suggest increased isotopic discrimination (fractionation) during photosynthesis, associated with greater reliance on passive CO₂ diffusion and/or internal CO₂ recycling as CCM efficiency declines under heat stress (80). In fact, all but one replicate in the 20-30°C range had δ¹³C < - 30, which falls into the third category described by Velázquez-Ochoa et al. 2022 (55), CO_2_ diffuse entry. The majority of replicates from the 5-15°C treatments had lower δ¹³C values, falling into category 2 (-10 < δ¹³C > -30), using a mix of CO_2_ and HCO_3_^-^ uptake (55). This mechanistic interpretation aligns with Bunting et al. 2024 (42), who demonstrated that marine heatwave intensity and duration suppress growth in young *M. pyrifera*, likely reducing CCM operation and tissue turnover, a process further supported here by elevated δ¹⁵N at thermal extremes, indicative of stress-limited nitrogen assimilation (42,82). Yet, compared to the performance experiment isotopic signatures, we found less negative δ¹³C and stable %C, %N, and C:N across treatments, indicating that priming did not substantially alter overall nutrient balance. These results do not necessarily confer priming buffers negative CCM impacts from heat stress, but rather there is no added burden and reductions in efficiency occurs at the most extreme temperatures (>20°C ). In all, marine heatwaves push kelp into a low-efficiency metabolic state by suppressing growth and carbon-concentrating mechanisms, which can undermine the resilience of kelp forests in the absences of buffering processes, that may include priming.

While our study was limited due to space and time with confounding factors arising from our two-phase approach, it was an important step forward for seaweed priming. We did find evidence of a ‘phase effect’ for several of the metrics (e.g., RGR, average *Fv/Fm*) that confounds some of our findings. While hatcheries have protocols in place to mitigate the chances of a bacterial or disease outbreak, this presents a real-world issue within seaweed and other aquaculture ventures (83,84). Although we did not see continued bacteria growth once phase 2 began at UC Santa Barbara, it is possible there were fewer sporophytes during this phase due to early impacts from the bacteria or possibly self-thinning – where small individuals crowd each other as they grow, and progressively die as competition increases (85) – as these blades were slightly larger than phase 1 blades. Increased *Fv/Fm*, although minimal between phases, could also be a result of differing age (86). Regardless, while the exact effect size of the fully versus just heat primed spools has higher uncertainty, the evidence is in favor of priming. Indeed, even with this phase effect, heat priming did not have a negative impact on any metric, underscoring that priming is still beneficial. For *M. pyrifera*, the next steps should focus on optimal duration of a priming, recovery, and timing of the trigger. Our trigger event was soon after the priming stress, but the best-case scenario for seaweed aquaculture practitioners would be if the priming stress contributed to longer-term (outplanting to harvest = 3-6 months) resilience against MHWs. Studies should also focus on epigenetic testing to understand the mechanisms underlying priming for seaweed species, as this has huge implication for the longevity of the priming effects (19). Ultimately, determining the best and most feasible protocols to ensure future resilience in a real-world setting will give aquaculture practitioners confidence in applying these techniques in their own hatcheries.

## CONCLUSIONS

Our findings suggest that adding a heat and nutrient priming intervention during the hatchery stage may improve outcomes for farms that experience marine heatwaves after outplanting. Importantly, the ‘high nutrient’ priming treatment we used in the study matches the industry standard for seaweed aquaculture, and thus adding a heat priming intervention would be relatively low effort and expense during the hatchery stage. As noted, this is a promising development as nutrient rescue has not been successful in *N. lutekeana* (67). While further studies should explore how long priming in *M. pyrifera* will result in heatwave protection and if other kelp species can be successfully primed, the applications for priming *M. pyrifera* could extend outside of aquaculture and into kelp conservation efforts.

## ACKNOWLEDGEMENTS

We thank Ocean Rainforest for their support and partnership in growing the early-stage sporophytes, with special thanks to Sam Reeves and Ann Bishop. We thank Shalanda Grier, Sara Matsumura, and Jayne Campbell for their logistical support breaking down this experiment.

