## Supplementary material for "Climate stress priming of juvenile Southern California giant kelp (*Macrocystis pyrifera*) to thermal extremes": SI tables & figs

PREPRINT

**S1 Table: Model selection for coverage proportion over time in the kelp performance experiment**

| Modnames | K | AICc | Delta_AICc | ModelLik | AICcWt | LL | Cum.Wt |
|---|---|---|---|---|---|---|---|
| day * treatment | 19.00 | -116.21 | 0.00 | 1.00 | 0.78 | 84.41 | 0.78 |
| day * treatment + level | 20.00 | -113.27 | 2.94 | 0.23 | 0.18 | 84.87 | 0.96 |
| day | 4.00 | -108.25 | 7.96 | 0.02 | 0.01 | 58.43 | 0.97 |
| intercept | 2.00 | -107.37 | 8.84 | 0.01 | 0.01 | 55.77 | 0.98 |
| day + treatment | 9.00 | -107.35 | 8.86 | 0.01 | 0.01 | 64.13 | 0.99 |
| treatment | 7.00 | -107.14 | 9.07 | 0.01 | 0.01 | 61.44 | 1.00 |
| day + treatment + level | 10.00 | -105.20 | 11.01 | 0.00 | 0.00 | 64.40 | 1.00 |
| day + treatment + jar | 27.00 | -69.09 | 47.12 | 0.00 | 0.00 | 78.73 | 1.00 |
| day * treatment + jar | 37.00 | -67.80 | 48.41 | 0.00 | 0.00 | 112.25 | 1.00 |

Note: Model selection was based on AICc. “Level” indicates the top of bottom of level of the experimental tanks and “jar” indicates the 3L jar used for each replicate.

**S2 Table: Summary output for the best fit model determined in S1 Table**

| *Predictors* | *Estimates* | *CI* | *p* |
|---|---|---|---|
| (Intercept) | 1.41 | 1.03 – 1.92 | **0.029** |
| day [7] | 0.91 | 0.59 – 1.40 | 0.657 |
| day [17] | 0.93 | 0.60 – 1.44 | 0.755 |
| treatment c [10°C] | 0.85 | 0.55 – 1.32 | 0.474 |
| treatment c [15°C] | 0.87 | 0.56 – 1.34 | 0.516 |
| treatment c [20°C] | 1.19 | 0.76 – 1.84 | 0.444 |
| treatment c [25°C] | 0.91 | 0.59 – 1.40 | 0.662 |
| treatment c [30°C] | 0.94 | 0.61 – 1.46 | 0.790 |
| day [7] × factor(treatment c)10°C | 1.00 | 0.54 – 1.84 | 0.993 |
| day [17] × factor(treatment c)10°C | 2.38 | 1.27 – 4.47 | **0.007** |
| day [7] × factor(treatment c)15°C | 1.29 | 0.70 – 2.38 | 0.418 |
| day [17] × factor(treatment c)15°C | 3.74 | 1.94 – 7.19 | **<0.001** |
| day [7] × factor(treatment c)20°C | 0.89 | 0.48 – 1.65 | 0.705 |
| day [17] × factor(treatment c)20°C | 1.36 | 0.72 – 2.54 | 0.340 |
| day [7] × factor(treatment c)25°C | 1.15 | 0.62 – 2.13 | 0.650 |
| day [17] × factor(treatment c)25°C | 0.85 | 0.46 – 1.58 | 0.613 |
| day [7] × factor(treatment c)30°C | 1.20 | 0.65 – 2.23 | 0.553 |
| day [17] × factor(treatment c)30°C | 0.68 | 0.37 – 1.26 | 0.223 |
| Observations | 72 | | |
| $R^2$ | 0.533 | | |

PREPRINT

**S3 Table: Model selection for coverage proportion over time in the kelp priming experiment**

| Modnames | K | AICc | Delta_AICc | ModelLik | AICcWt | LL | Cum.Wt |
|---|---|---|---|---|---|---|---|
| day * nutrient * prime | 13.00 | -240.02 | 0.00 | 1.00 | 0.54 | 134.41 | 0.54 |
| day * nutrient * prime + level | 14.00 | -238.35 | 1.67 | 0.43 | 0.23 | 134.80 | 0.78 |
| day * nutrient + trial | 8.00 | -238.08 | 1.95 | 0.38 | 0.20 | 127.57 | 0.98 |
| day + prime + trial | 6.00 | -233.07 | 6.95 | 0.03 | 0.02 | 122.84 | 1.00 |
| day + nutrient + prime | 6.00 | -228.40 | 11.63 | 0.00 | 0.00 | 120.50 | 1.00 |
| day + nutrient + prime + level | 7.00 | -227.37 | 12.66 | 0.00 | 0.00 | 121.09 | 1.00 |
| day + nutrient | 5.00 | -225.74 | 14.29 | 0.00 | 0.00 | 118.09 | 1.00 |
| day + nutrient + level | 6.00 | -224.72 | 15.31 | 0.00 | 0.00 | 118.67 | 1.00 |
| day * nutrient | 7.00 | -223.39 | 16.63 | 0.00 | 0.00 | 119.11 | 1.00 |
| day * nutrient + level | 8.00 | -222.36 | 17.66 | 0.00 | 0.00 | 119.71 | 1.00 |
| day + prime | 5.00 | -219.22 | 20.80 | 0.00 | 0.00 | 114.83 | 1.00 |
| day + prime + level | 6.00 | -218.27 | 21.75 | 0.00 | 0.00 | 115.44 | 1.00 |
| day | 4.00 | -217.98 | 22.04 | 0.00 | 0.00 | 113.13 | 1.00 |
| day * prime | 7.00 | -216.40 | 23.62 | 0.00 | 0.00 | 115.61 | 1.00 |
| day * prime + level | 8.00 | -215.39 | 24.64 | 0.00 | 0.00 | 116.23 | 1.00 |
| day * nutrient * prime + jar | 57.00 | -204.08 | 35.95 | 0.00 | 0.00 | 197.48 | 1.00 |

| Modnames | K | AICc | Delta_AICc | ModelLik | AICcWt | LL | Cum.Wt |
|---|---|---|---|---|---|---|---|
| day + nutrient + jar | 51.00 | -200.99 | 39.04 | 0.00 | 0.00 | 180.32 | 1.00 |
| day + prime + jar | 51.00 | -200.99 | 39.04 | 0.00 | 0.00 | 180.32 | 1.00 |
| day * prime + jar | 53.00 | -195.18 | 44.84 | 0.00 | 0.00 | 182.39 | 1.00 |
| day + nutrient + prime + jar | 53.00 | -195.13 | 44.89 | 0.00 | 0.00 | 182.37 | 1.00 |
| day * nutrient + jar | 53.00 | -195.13 | 44.89 | 0.00 | 0.00 | 182.37 | 1.00 |
| nutrient + prime + trial | 5.00 | -168.25 | 71.77 | 0.00 | 0.00 | 89.34 | 1.00 |
| nutrient * prime | 5.00 | -165.43 | 74.60 | 0.00 | 0.00 | 87.93 | 1.00 |
| nutrient * prime + level | 6.00 | -163.80 | 76.22 | 0.00 | 0.00 | 88.21 | 1.00 |
| nutrient + prime | 4.00 | -161.17 | 78.85 | 0.00 | 0.00 | 84.73 | 1.00 |
| nutrient | 3.00 | -160.32 | 79.70 | 0.00 | 0.00 | 83.25 | 1.00 |
| nutrient + prime + level | 5.00 | -159.79 | 80.23 | 0.00 | 0.00 | 85.11 | 1.00 |
| intercept | 2.00 | -156.54 | 83.49 | 0.00 | 0.00 | 80.31 | 1.00 |
| prime | 3.00 | -156.46 | 83.56 | 0.00 | 0.00 | 81.32 | 1.00 |
| nutrient * prime + jar | 49.00 | -79.14 | 160.88 | 0.00 | 0.00 | 114.63 | 1.00 |
| nutrient + prime + jar | 49.00 | -79.14 | 160.88 | 0.00 | 0.00 | 114.63 | 1.00 |

Note: Model selection was based on AICc. “Level” indicates the top of bottom of level of the experimental tanks, “jar” indicates the 3L jar used for each replicate, and “trial” represents the phase of the experiment.

**S4 Table: Summary output for the best fit model determined in S2 Table**

| *Predictors* | *Estimates* | *CI* | *p* |
|---|---|---|---|
| (Intercept) | 0.41 | 0.32 – 0.53 | **<0.001** |
| day [7] | 1.29 | 0.90 – 1.84 | 0.167 |
| day [17] | 2.53 | 1.78 – 3.59 | **<0.001** |
| nutrient id [full] | 1.15 | 0.81 – 1.62 | 0.441 |
| prime id [primed] | 1.14 | 0.81 – 1.62 | 0.454 |
| day [7] × factor(nutrient id)full | 0.92 | 0.57 – 1.50 | 0.750 |
| day [17] × factor(nutrient id)full | 0.74 | 0.46 – 1.19 | 0.212 |
| day [7] × factor(prime id)primed | 0.78 | 0.48 – 1.27 | 0.314 |
| day [17] × factor(prime id)primed | 0.67 | 0.42 – 1.08 | 0.103 |
| nutrient id [full] × factor(prime id)primed | 0.97 | 0.60 – 1.58 | 0.907 |
| day [7] × factor(nutrient id)full × factor(prime id)primed | 1.49 | 0.75 – 2.93 | 0.253 |
| day [17] × factor(nutrient id)full × factor(prime id)primed | 3.23 | 1.65 – 6.33 | **0.001** |
| Observations | 144 | | |
| $R^2$ | 0.502 | | |

PREPRINT

**S1 Figure: Temperature differences across kelp performance experiment**

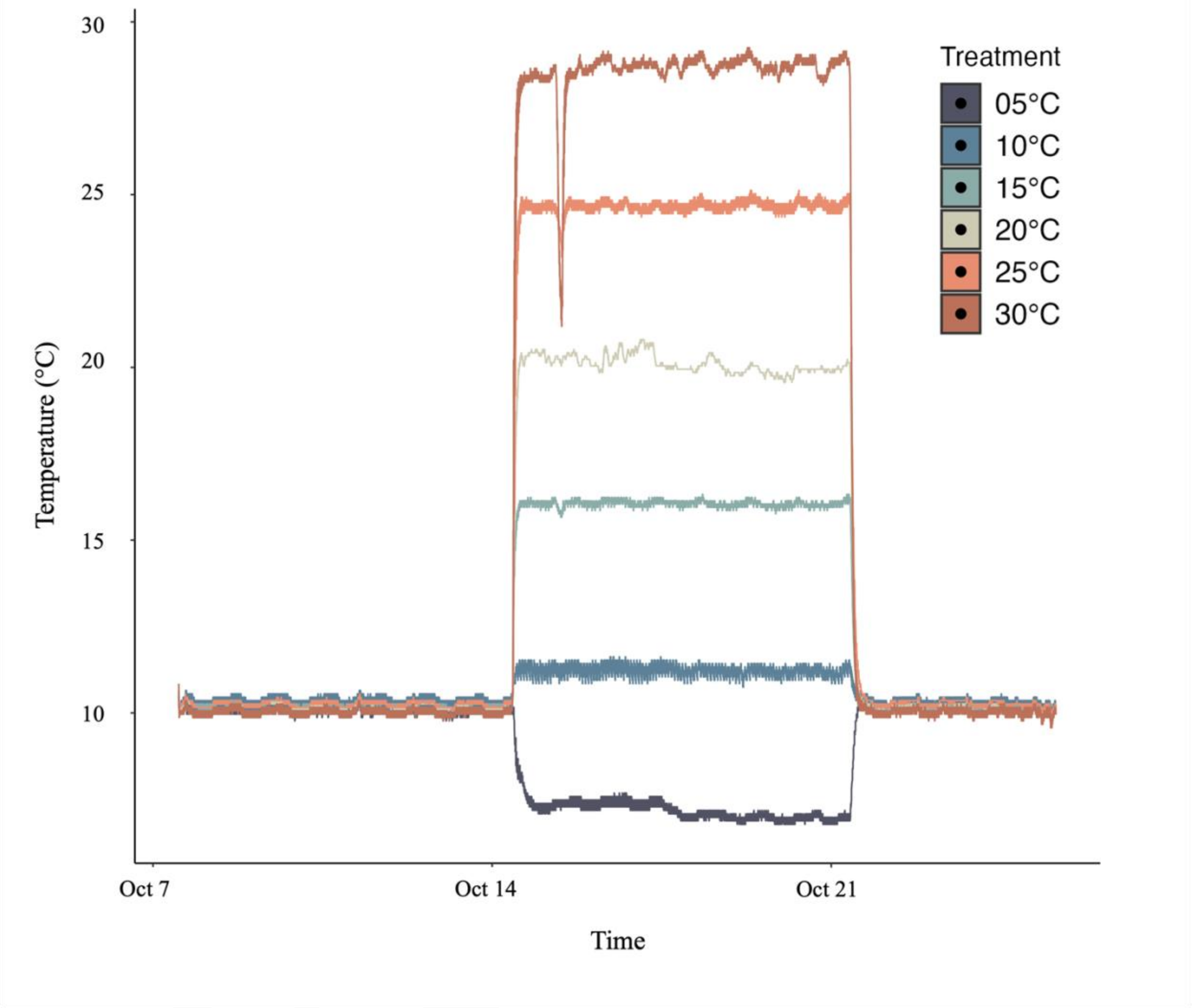


Note. Onset HOBO logger temperature readings over the kelp performance experiment.

PR

**S2 Figure: Temperature across phase 1 of the kelp priming experiment**

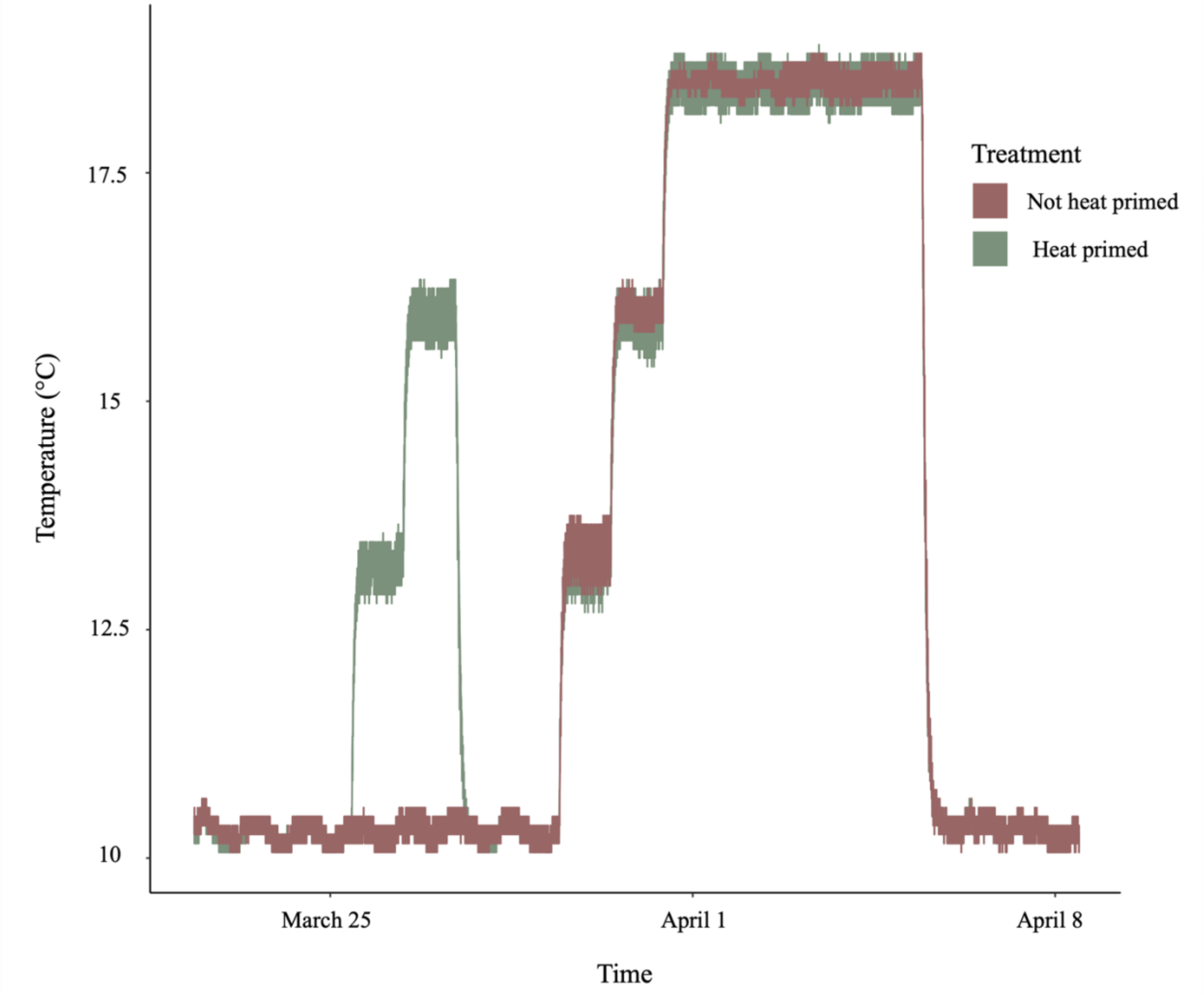


Note. Onset HOBO logger temperature readings for phase 1 of the kelp priming experiment.

PR

**S3 Figure: Temperature across phase 2 of the kelp priming experiment**

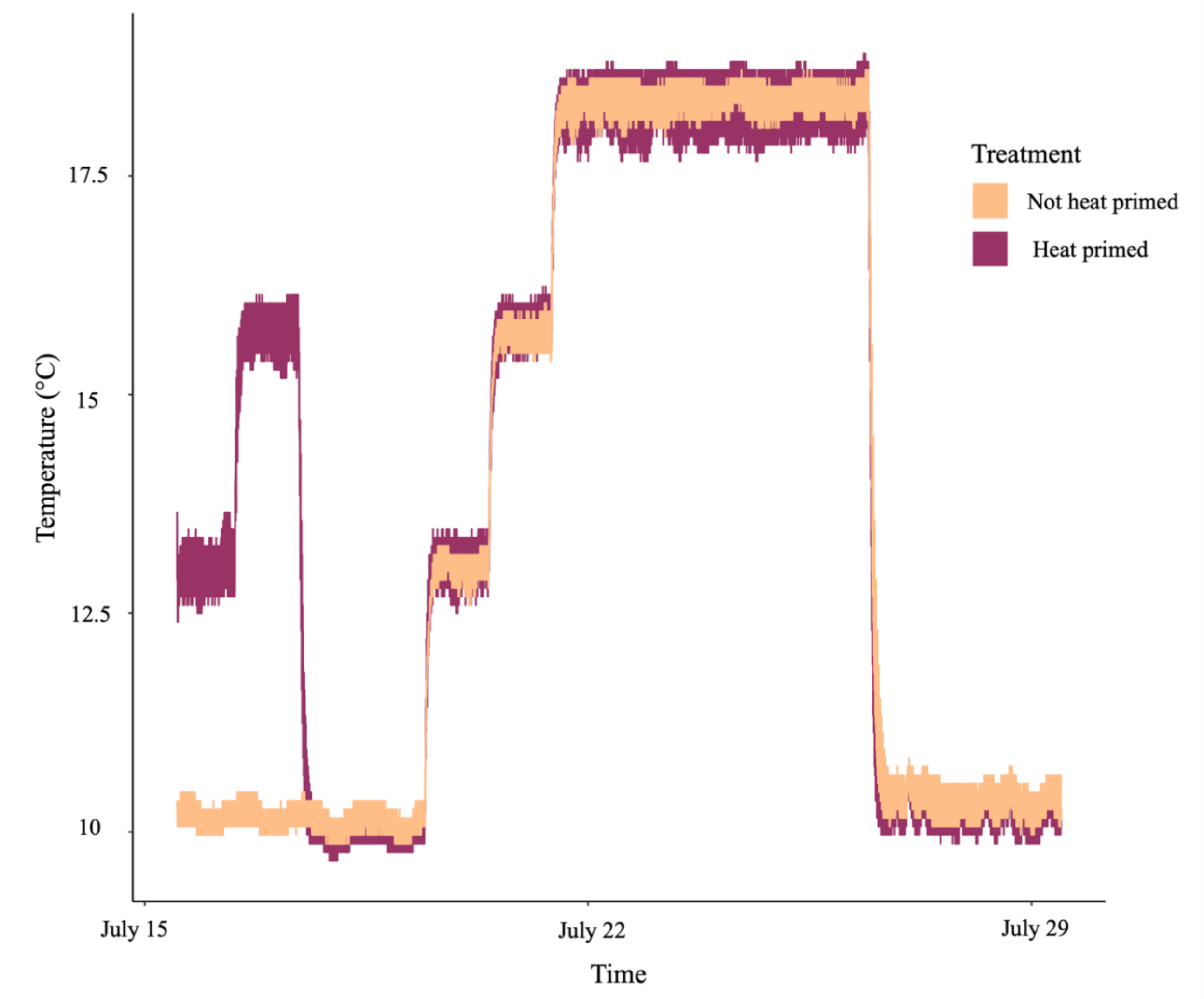


Note. Onset HOBO logger temperature readings for phase 2 of the kelp priming experiment.

Loggers began recording at the start of the heat priming.

PR

**S4 Figure: Coverage proportion over time for the kelp performance experiment**

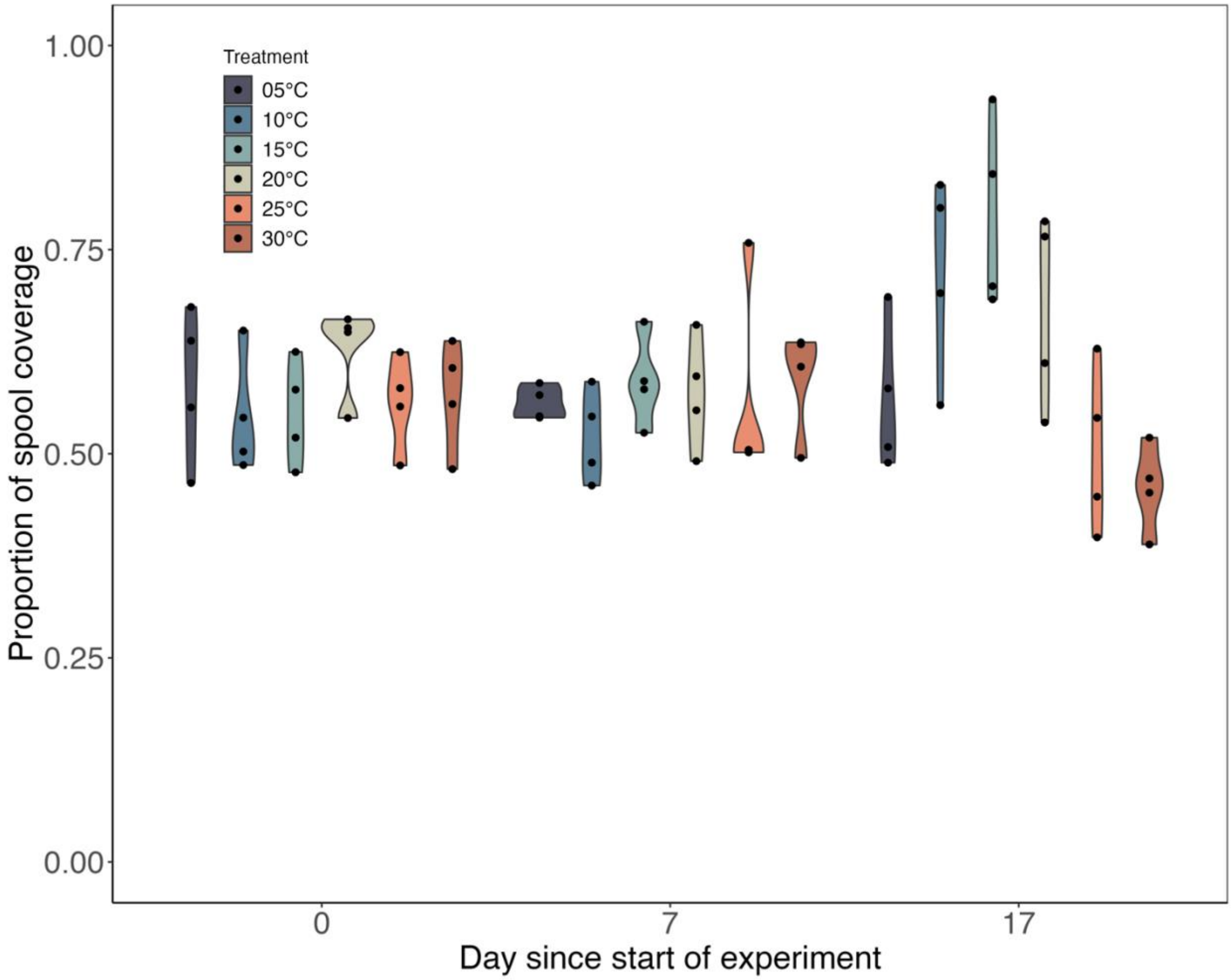


PRL

Note. Proportion of mini-spool coverage at the start of the kelp performance experiment (Day 0), after the hatchery stage (Day 7), and at the conclusion of the experiment (Day 17).

**S5 Figure: Cluster analysis on nitrogen and carbon relationship in the kelp performance experiment**

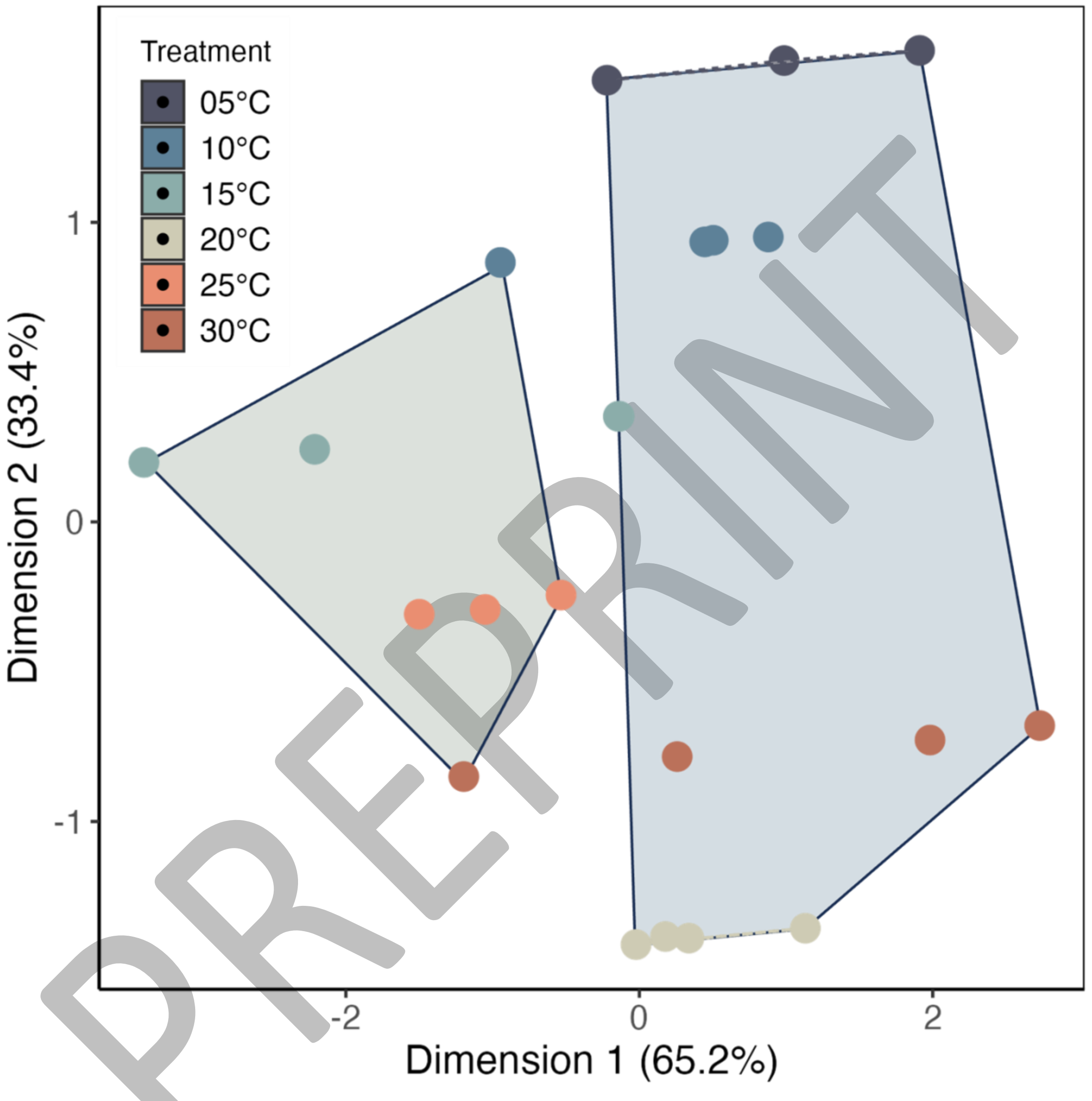


Note. Cluster analysis to examine the carbon and nitrogen relationship and temperature. Extreme heat does not stand out.
